# Spontaneous neurotransmitter release is regulated by Unc-5

**DOI:** 10.64898/2026.06.02.729526

**Authors:** Samuel W. Vernon, Evelyne Ruchti, Chiara Paolantoni, Rebecca C. Smith, Marine Van Campenhoudt, Brian D. McCabe

## Abstract

The spontaneous quantal release of single vesicles of neurotransmitter in the absence of an action potential is a universal feature of all neuronal chemical synapses. These ‘miniature events’ have long been thought to be stochastic and unregulated, leading to measurement of their frequency as a widely employed estimate of the number of synaptic connections within neuronal networks. Here we show using high resolution live imaging of *Drosophila* adult glutamatergic synapses that the spontaneous release of neurotransmitter occurs from only a subset of synaptic release sites. We discover that the proportion of release sites participating in spontaneous neurotransmitter release is regulated by a novel synaptic function of the transmembrane signalling receptor Unc-5 which is heparan sulfate proteoglycan dependent, but Netrin independent. We show that synaptic Unc-5 forms a complex with the SNARE protein Syntaxin to regulate neurotransmitter release. The depletion of Unc-5 in adult synapses diminishes miniature events, inducing terminal degeneration and behavioural decline. Our results reveal that ‘spontaneous’ neurotransmitter release is a singular, essential and independently regulated property of synapses that is critical for the structural integrity and behavioural contribution of synaptic connections.

## Introduction

Individual vesicular packets of neurotransmitter, or quanta, are released at a defined rate (Probability of release — *P*_*r*_) from specialised release sites, known as active zones (AZs) within synaptic terminals, either co-ordinately, in response to electrical stimulation during evoked neurotransmitter release, or individually in the absence of stimulation, known as spontaneous neurotransmitter release or alternatively miniature neurotransmission. Miniature neurotransmission was first described over 75 years ago(1, 2) and, together with subsequent studies(3–7), led to the establishment of the quantal mechanism of vesicular release which underpins our current understanding of neurotransmission at every fast chemical synapse. Implicit in this model was the assumption that active zones are functionally homogenous and that all active zones produce both evoked and spontaneous vesicular release events(8–10). These foundational studies operated under the assumption that spontaneous vesicular release is an unregulated epiphenomena or noise(11). Based on these two assumptions, the measurement of the frequency of miniature events was used as an electrophysiological proxy for the number of synaptic connections a neuron receives. However, recent technical advances have begun to question some of these core presumptions. First, though correlation between the number of synaptic structures and miniature event frequency has been observed in some systems(12–17), numerous other studies have demonstrated a disconnect between the number of synapses and miniature event frequency(18–21). Second, cumulative evidence has established that baseline spontaneous and evoked neurotransmission can be functionally and molecularly segregated at rest(22–33), and also display a capacity for stimulation dependent plasticity(27, 31). Finally, the view that spontaneous vesicular release is merely an epiphenomena has been questioned by studies showing distinct functional requirements for miniature neurotransmission including for synapse development and maintenance(34–39).

Here we further challenge a central pillar of prevailing models of neurotransmission by showing that spontaneous release is, in fact, a regulated process restricted to specialised release sites with dedicated molecular machinery that govern it as an independent neurotransmission modality. Using optical methods to examine the spatial properties of evoked and spontaneous release events with single AZ spatial resolution in mature adult *Drosophila* glutamatergic synapses, we observe AZ segregation of evoked and miniature neurotransmission. From a genetic screen, we discover that Unc-5, a transmembrane signalling receptor, best known for its role as a Netrin-dependent axon guidance receptor, is enriched at synaptic AZs that release miniature events. We find that increasing or decreasing synaptic Unc-5 levels can increase or decrease spontaneous release frequency. We further show that the regulation of spontaneous neurotransmitter release events by Unc-5 is critical to maintain adult synaptic structural integrity and behavioural capacity. Our results establish that spontaneous neurotransmission is not only segregated from, but also independently regulated from, evoked neurotransmitter release, and that spontaneous events have a singular and essential role for structural maintenance of adult synaptic connections and behaviour.

## Results

### Evoked and spontaneous neurotransmitter release is segregated at adult synapses

We recently developed a mature adult *Drosophila* glutamatergic neuromuscular synapse preparation which enables genetic, electrophysiological and imaging access throughout adult life(38, 39) (Fig. 1a). Active zones are specialised regions of synapses where presynaptic neurotransmitter vesicular release components are closely opposed to postsynaptic neurotransmitter receptors(40) and we wished to visualise neurotransmitter release with individual AZ spatial resolution in these terminals.

**Figure 1.**
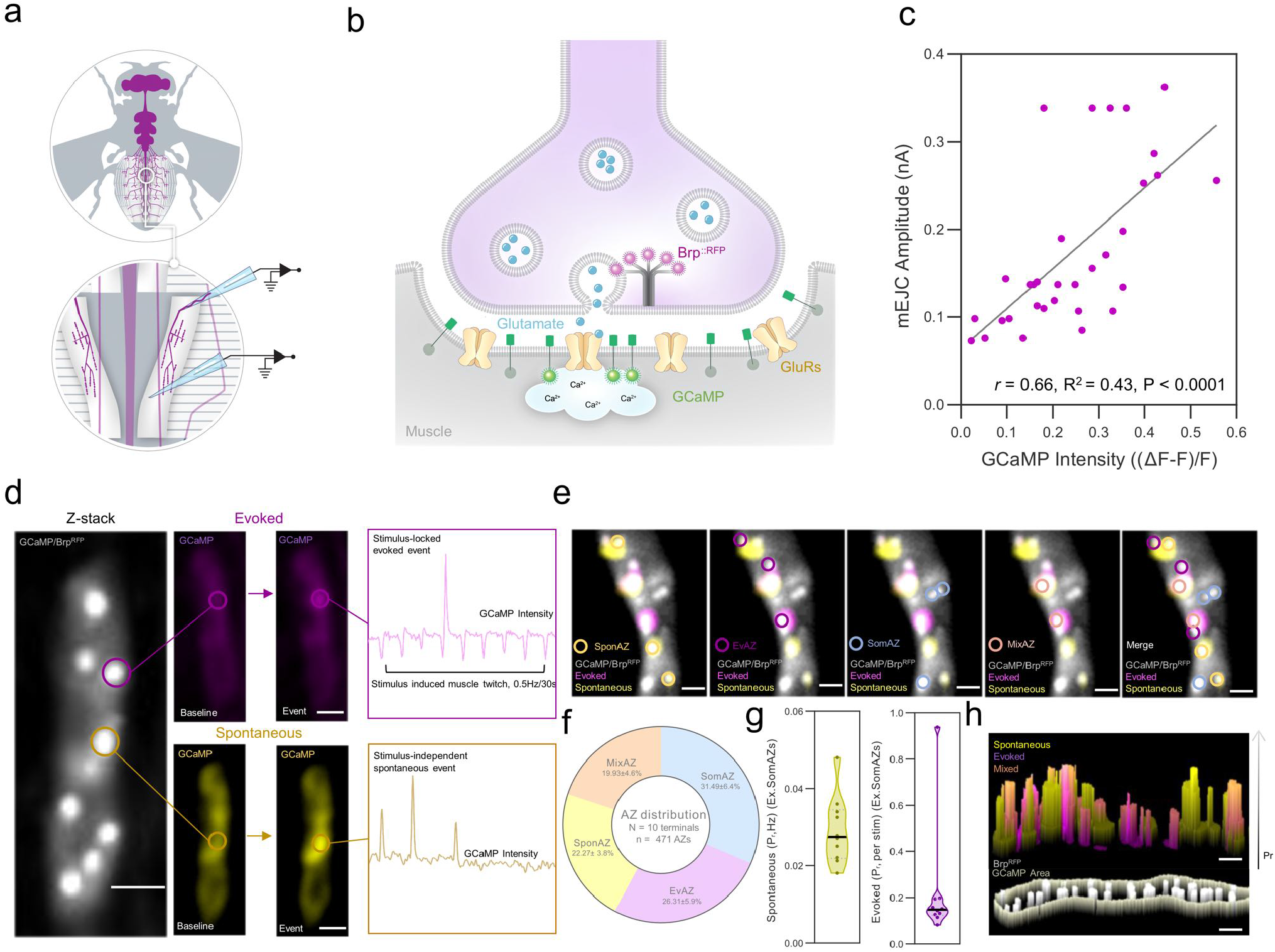
Adult *Drosophila* NMJ terminals display spatial segregation of release modalities. **a**, Schematic of targeted adult *Drosophila* abdominal MVIM abdominal muscles. **b**, Schematic of live imaging strategy. Endogenously tagged Brp::RFP (magenta) together with postsynaptic localised GCaMP (SynapGCaMP6f, green) allow functional live imaging at individual AZs without a requirement for post-hoc immunohistochemistry. Quantal release of glutamate (blue) triggers excitatory ligand-gated GluR (yellow) activation visualised through a corresponding localised Ca2+ mediated flash of GCaMP (green) localised to the postsynaptic SSR. **c**, Recorded GCaMP optical events displayed a 1:1 concurrence with recorded mEJCs within the same fibre with GCaMP amplitudes showing a modest positive linear correlation with mEJC amplitudes (*r* = 0.66, *P <* 0.0001, *R*^2^ = 0.43, *P <* 0.0001, *n* = 32). **d**, Schematic demonstrating spatial and temporal identification of spontaneous (yellow) and evoked (magenta) postsynaptic Ca2+ events (see methods for further details, Scale Bars: 3 µm). **e**, AZs were characterised as spontaneous (SponAZ, yellow), evoked (EvAZ, magenta), silent/somnolent (SomAZ, blue) or mixed (MixAZ, pink) based on the presence or absence of specified release modality during recording period. **f**, Quantification of AZ subtype spatial distribution reveals functional heterogeneity at individual release sites. Analysis was conducted on both spatial distribution and temporal probability of release quantified here in **g**, violin plots for mean spontaneous *Pr* (yellow) and mean evoked *Pr* (magenta) per recording (excluding SomAZs) and represented in **h**, as a 3D cumulative count event map utilised for visualisation purposes only (*n* = 10 recordings, scale bars: 2 µm). All statistical analysis details are reported in Supplementary Tables and in Source Data file. Violin Plots represent median & IQR, distribution data is presented as mean *±* SEM.

To do this we used genome engineering to introduce mScarlet(41) in frame into the presynaptic AZ protein Bruchpilot (Brp)(42). We coupled this presynaptic AZ marker with postsynaptic localised GCaMP(27) (Fig. 1b) enabling us to simultaneously image evoked and spontaneous vesicular release events at individual AZs with single vesicle resolution over a time period of up to 30 minutes (Fig. 1d–e). To ensure this imaging system accurately captured neurotransmission events, we also simultaneously carried out electrophysiological recordings from postsynaptic muscles (Fig. S1a–b).

To evaluate the system, we began by establishing that the postsynaptic calcium imaging events we observed were directly attributable to presynaptic vesicular neurotransmitter release. First, we silenced all presynaptic vesicular release (both evoked and spontaneous) with conditional adult expression of transgenic botulinum toxin (BoNT)(43) and, as expected found that the frequency of postsynaptic calcium events was dramatically reduced (Fig. S1c–j). Second, we increased spontaneous and evoked vesicular neurotransmission by genetic depletion of the vesicular release clamp Tomosyn(44) and saw a large increase in the frequency of postsynaptic calcium events (Fig. S1c–e). Finally, we confirmed that vesicular fusion events predicted by functional imaging were coincident with the expected postsynaptic electrophysiological measurements with one-to-one concordance (Fig. 1c, Fig. S1a–b). These results confirmed that postsynaptic calcium imaging using these tools accurately reflected presynaptic vesicular release events with spatial and temporal precision.

Having established the fidelity of our imaging system, we next examined the spatial distribution of both evoked and spontaneous neurotransmitter release at individual AZs. We observed that the release properties of individual AZs within mature adult terminals were heterogeneous (*N* = 10 terminals, *n* = 471 AZs, Fig. 1e–f). We found that when neurons were stimulated at 0.5 Hz, 19.93 *±* 4.63% of AZs produced both spontaneous and evoked events. However, of the remaining AZs, 26.31 *±* 5.92% of AZs produced only evoked events and a further 22.27 *±* 3.82% produced only spontaneous vesicular release events. Finally, 31.49 *±* 6.43% of AZs were observed to not release vesicles at all during evoked stimulation and also not to participate in spontaneous release during the imaging period (Fig. 1f). We examined this last population of AZs more closely. We found that these AZs could elicit postsynaptic responses upon focal application of sucrose, KCl or glutamate (Supplemental Movie 1a–c) indicating that they were capable of neurotransmitter release. Supporting this, through post-hoc immunolabeling, we found these AZs had opposing postsynaptic glutamate receptors (Supplemental Movie 1d–e). Given that these AZs appeared functional but did not produce vesicular release events under our stimulation conditions, we dubbed these quiescent AZs ‘somnolent’. In sum, our imaging found four categories of vesicular release AZs: those with a high probability of release during evoked neurotransmission but a low probability of spontaneous release (EvAZ), AZs which participate in both forms of neurotransmission (MixAZ), AZs which produce spontaneous events primarily (SponAZ) and somnolent AZs with a low probability of either form of vesicular release (SomAZ). Overall, adult *Drosophila* terminals contain release sites with divergent average probabilities of spontaneous and evoked release (Fig. 1g–h) and spatially heterogeneous release modalities.

### Unc-5 modulates active zone release probabilities

We have previously found that the specific inhibition of spontaneous neurotransmission induces a characteristic structural degeneration of adult synaptic terminals, in particular the subdivision or fragmentation of synaptic boutons(38, 39). Leveraging this morphological phenotype, we screened for proteins which, when selectively inhibited in adult neurons using temporally controlled RNA interference (RNAi), could alter spontaneous neurotransmission. From this effort, we identified a novel role for the transmembrane signalling receptor Unc-5. Unc-5 has multiple physiological roles(45–47) but is perhaps best known as a receptor for Netrin during neuronal axon guidance(45).

When we conditionally inhibited Unc-5 in adult motor neurons (MNs) using RNAi we observed a 60.98% (*P <* 0.001) decrease in the frequency of spontaneous neurotransmission compared to controls without any change in the amplitude of these events (Fig. 2a–c). This was accompanied by a 68.79% (*P <* 0.001) decrease in excitatory junctional current (EJC) amplitudes and a 72.9% (*P <* 0.001) decrease in quantal content (*P <* 0.001), a measure of the number of vesicles released per stimulus, compared to controls (Fig. 2d–e). We confirmed this result was specific by nanobody based degradation(48) of Unc-5 protein in adult synapses, where we also observed a 63.22% (*P <* 0.05) decrease in the frequency of spontaneous neurotransmission also without changing the amplitude of these events, and similarly this was accompanied by a 33.33% decrease in EJC amplitudes with a 35.83% decrease in quantal content (*P <* 0.05) (Fig. S2a–e). Intriguingly, we found that both the decrease of EJC amplitudes and the quantal content decline we observed when Unc-5 was depleted could be rescued by elevating extracellular (Fig. S2f–i) or intracellular (Fig. S2l–r) Ca^2+^ availability. In contrast, the decline of miniature excitatory junctional current (mEJC) frequency when Unc-5 was depleted was not rescued by Ca^2+^ manipulation (Fig. S2j–k,s–t). This disassociation suggested that the depletion of Unc-5 impaired evoked and spontaneous neurotransmission through distinct mechanisms, with effects upon spontaneous event frequency not explained by a shared Ca^2+^-dependant process. We next examined the effects of overexpressing Unc-5 in adult motor neurons. We found that, opposite to Unc-5 depletion, increasing Unc-5 levels induced a 157.2% (*P <* 0.001) increase in the frequency of spontaneous neurotransmission events, again without affecting their amplitudes (Fig. 2a–c). Strikingly, when Unc-5 was overproduced, we observed no significant increase in evoked amplitudes and no change in quantal content (Fig. 2d–e). This result again suggested that Unc-5 regulates evoked and spontaneous release through distinct mechanisms, and further revealed that the levels of Unc-5 can strongly and bidirectionally influence spontaneous neurotransmission frequency.

**Figure 2.**
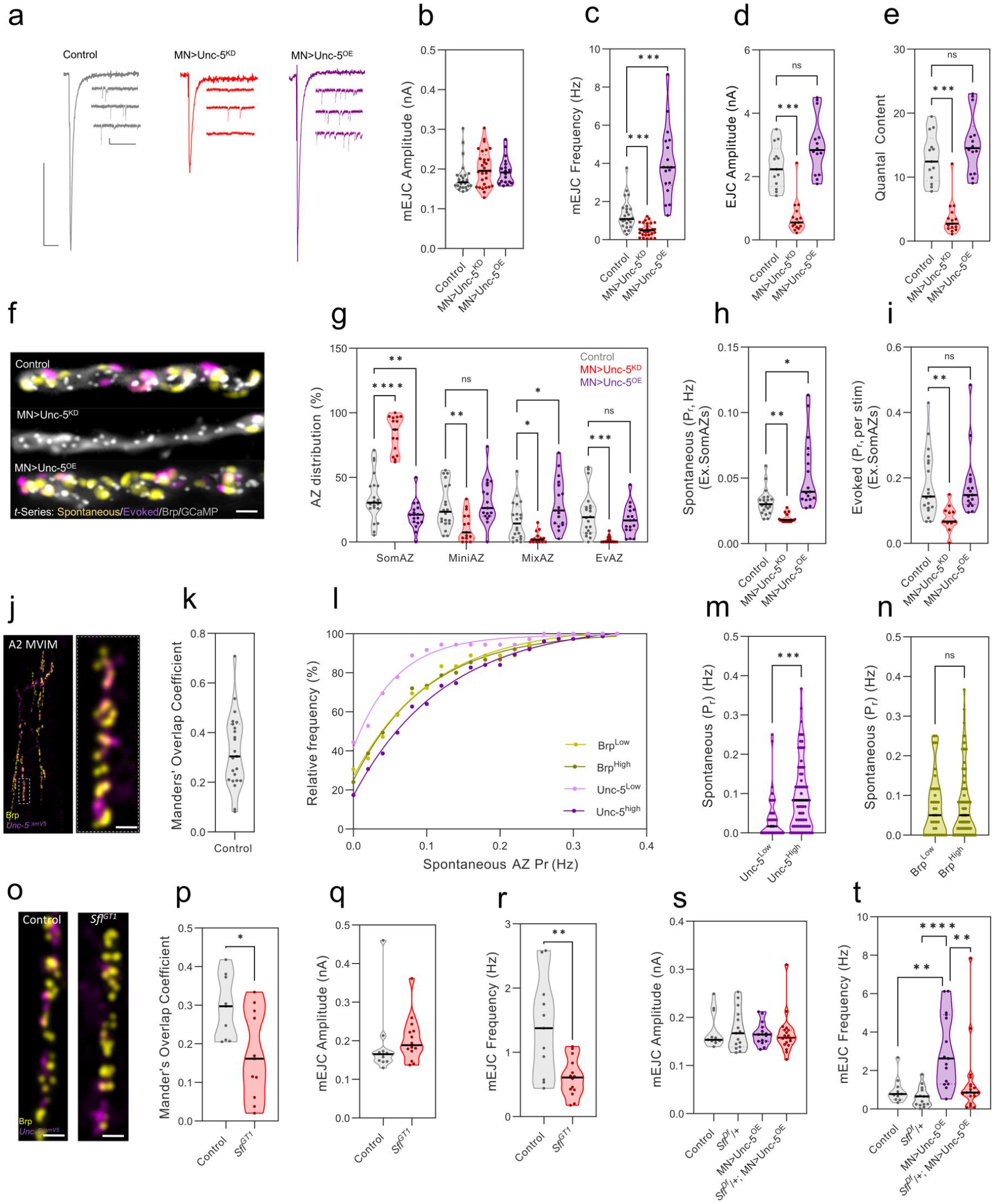
Unc-5 localises to subsets of AZs and bidirectionally influences spontaneous neurotransmitter release. **a**, Representative traces of excitatory glutamatergic currents (Scale Bars: EJCs, 1 nA/30 ms mEJCs, 0.1 nA/500 ms). **b–e**, Violin plots showing Unc-5 expressional dysregulation does not influence mEJC amplitudes, bidirectionally influences mEJC frequency (*n >* 18 recordings per genotype, Kruskal-Wallis test: *P <* 0.0001) and unidirectionally reduces EJC amplitude and quantal content (*n <* 13 recordings per genotype, Kruskal-Wallis test: *P <* 0.0001). **f**, Representative *t*-Series stack showing spontaneous (yellow) and evoked (magenta) activity maps together with Brp/GCaMP (grey) localisation for Control, MN*>*Unc-5KD and MN*>*Unc-5OE (Scale Bar: 3 µm). **g**, Violin plots showing Unc-5 levels correlate with the functional distribution of AZ activity observed during live imaging (*n >* 15 recordings per genotype, 2-Way ANOVA: *P <* 0.0001) reflected in **h**, bidirectional modulation of spontaneous *Pr* at active individual release sites (*n >* 15 recordings per genotype ex.SomAZs, Kruskal-Wallis test: *P <* 0.0001) together with **i**, unidirectional loss of function in Evoked *Pr* at active release sites (*n >* 15 recordings per genotype ex.SomAZs, Kruskal-Wallis test: *P <* 0.0001). **j**, Representative image of endogenously tagged Unc-5::smV5 (magenta) and AZ marker Brp (yellow) at full MVIM A2 terminal (left panel, Scale bar: 10 µm) together with high magnification subsection (right panel, Scale Bar: 3 µm). **k**, Violin plot showing partial colocalization of Unc-5 with AZs (*n* = 23 NMJs). **l**, Spontaneous cumulative frequency is left shifted at AZs occupied by relatively lower levels of Unc-5 (*<*30% relative IHC intensity, Unc-5Low, pink) relative to AZs occupied by relatively higher levels (*>*30% relative IHC intensity, Unc-5High, magenta). This trend was not observed for Brp levels (yellow) and is directly compared in violin plots **m–n**, (Violin Plots: *m, n* = 111 AZs, Mann Whitney test: *P <* 0.001, *n, P* = 0.75). **o**, Representative images of Unc-5::smV5 (magenta) and Brp (yellow) in control and Sfl hypomorph *sfl*GT1. **p**, *Sfl*GT1 induced a reduction in Unc-5 overlapping with Brp (Violin Plot: *n >* 9 NMJs per genotype, Student’s *t*-test, *P <* 0.05), correlative with a decline in mEJC frequency (Violin Plots: **q**, mEJC amplitude, *P* = 0.09, **r**, mEJC frequency, *n >* 11 recordings per genotype, Mann Whitney test: *P <* 0.01). **s**,**t**, The incorporation of a heterozygous deletion of *sfl* was sufficient to reduce MN*>*Unc-5OE mEJC frequency (*n >* 11 recordings per genotype, Kruskal-Wallis test, *P <* 0.001). Violin Plots represent median & IQR. All detailed statistical analysis & genotype details are reported in Supplementary Table 1

We next employed our spatial neurotransmission imaging system to examine how increasing or decreasing Unc-5 expression alters neurotransmitter release with active zone resolution. We observed that when Unc-5 expression was reduced in adult neurons, and as described above, the frequency of spontaneous vesicle release was reduced, the proportion of somnolent AZs was increased by 142.55% (*P <* 0.0001) (Fig. 2f–g). Concomitantly, the proportion of AZs that produced either spontaneous events alone ( *−*56.8%, *P <* 0.01), evoked events alone ( *−*92.2%, *P <* 0.001) or both forms of neurotransmission ( *−*79.8%, *P <* 0.05) was reduced (Fig. 2g). We then examined the release rate at individual AZs using our imaging system. We found that the probability of release at AZs that still produce spontaneous release was reduced ( *−*37.8%, *P <* 0.01) as was the rate at AZs that produced evoked neurotransmission ( *−*56.7%, *P <* 0.01) (Fig. 2h–i). We next examined what occurs at AZs when Unc-5 expression is increased. We observed that increasing Unc-5 decreased the percentage of somnolent AZs by 40.10% (*P <* 0.01). In parallel increasing levels of Unc-5 increased the proportion of MixAZs producing both spontaneous and evoked release (+81.5%, *P <* 0.05). The rate of spontaneous release at individual AZs was also increased (+69.2%, *P <* 0.05). In contrast, evoked active zone probability of release was unchanged (Fig. 2h–i). This spatial imaging data agreed with our electrophysiological results and showed that reducing or increasing the levels of Unc-5 in adult synapses changes the distribution and the rate of neurotransmitter release at individual AZs, with profound and bidirectional effects on AZs that produce spontaneous release independently of evoked release.

### Unc-5 is enriched at active zones participating in spontaneous release

We next used genome engineering(49) to insert an epitope tag, smFP-V5(50), in frame in the Unc-5 locus (Fig. S3a). We confirmed that this modified protein migrated at the expected molecular weight (Fig. S3b) and did not perturb Unc-5 function as the tagged protein was produced in a pattern similar to previous observations of Unc-5 expression during CNS development(51) (Fig. S3c), rescued the lethality of Unc-5 mutants (Fig. S3d), and did not alter neurotransmission (Fig. S3e–i). We then used this tagged allele to examine the expression of Unc-5 in adult synapses. We found that Unc-5 was localised to a subset of AZs (Fig. 2j–k), which we quantified as 42.75% of the total amount of AZs (*N* = 14 terminals, *n* = 3739 AZs). We then used our spatial neurotransmission imaging system to examine the release properties of AZs that had high or low levels of Unc-5. We observed that AZs with low Unc-5 levels had a low probability of spontaneous release (Fig. 2l–m, *P <* 0.001). This relationship was not observed for other synaptic proteins such as Brp (Fig. 2n). In contrast, when we examined AZs enriched for Unc-5, we observed a positive linear correlation between Unc-5 levels and spontaneous release probability (*r* = 0.54, *P <* 0.0001, *R*^2^ = 0.29, *P <* 0.0001, Fig. S3j). Again, no such correlation was observed for Brp (Fig. S3k). Thus, in adult synapses, Unc-5 is enriched at a subset of AZs and these release sites have a higher probability of spontaneous neurotransmitter release.

### HSPGs, but not Netrin, are required for Unc-5 synaptic functions

In axon guidance and other contexts Netrin acts as a ligand for Unc-5(52, 53). The *Drosophila* genome contains two Netrin encoding genes, NetA and NetB, and loss-of-function double mutants of both genes can survive to adulthood(54). We examined adult Net^*AB*^ double mutants and found that abdominal muscles were innervated correctly and that terminals looked morphologically normal. Electrophysiological recordings from Net^*AB*^ double mutants revealed no alterations of either evoked or spontaneous neurotransmission (Fig. S4a–e). Furthermore, Unc-5 mediated changes of spontaneous vesicle release frequency were not altered when Netrin expression was depleted (Fig. S4e). In sum, these data did not support a role for Netrins in the regulation of neurotransmission by Unc-5.

In addition to Netrin, Heparan sulfate proteoglycans (HSPGs) have also been implicated in regulating Unc-5-dependent axon guidance(55, 56). HSPGs act through covalently attached sugar side chains that carry multiple structural modifications, including importantly, sulfate residues. The *sulfateless (sfl)* gene encodes the *Drosophila* heparan sulfate-glucosamine N-sulfotransferase which catalyses an essential first step in the process of sulfotransferase modifications of HSPGs(57). We found Sfl was localised to adult synapses (Fig. S4f). We next established that a hypomorphic *sfl*^*GT* 1^ loss-of-function mutant allele(58, 59) can survive until adulthood enabling us to examine Unc-5 levels and localisation in these mutants. We found that the total number of all AZs labelled by Unc-5 was reduced by 42.1% (*P <* 0.05) in *sfl* ^*GT* 1^ mutant synapses compared to controls, indicating that HSPG sulfation was required for Unc-5 localisation at synapses (Fig. 2o–p). We then examined the neurotransmission properties of *sfl*^*GT* 1^ mutants. We found they had a 56.19% (*P <* 0.01) reduction in spontaneous neurotransmitter release (Fig. 2q–r) frequency, consistent with the reduction we observed in the synaptic levels of Unc-5. We next tested if the increase of spontaneous neurotransmission frequency induced by Unc-5 overexpression was also dependent upon Sfl function. To do this, we overexpressed Unc-5 in adult neurons while simultaneously reducing Sfl levels through heterozygous genetic deletion of *sfl*. Unlike homozygous *sfl*^*GT* 1^ alleles, heterozygous deletion of *sfl* alone did not significantly reduce spontaneous neurotransmitter release (Fig. 2t). However, removal of one copy of *sfl* reduced the Unc-5 mediated increase in spontaneous release frequency by 54.8% (*P <* 0.001, Fig. 2s–t). This genetic interaction, together with immunolabeling data, supports that activity of Unc-5 upon neurotransmission is dependent upon Sfl levels and by extension synaptic HSPG function.

### Unc-5 modulates spontaneous release through its association with Syntaxin

As depletion of synaptic Unc-5 decreased spontaneous neurotransmission, we wondered if Unc-5 might interact with vesicular release ‘clamp’ proteins which also act to inhibit vesicular release(60). These proteins, which include Tomosyn and Complexin, increase the frequency of spontaneous neurotransmitter release when reduced with diverse effects on evoked release(44, 61). We first tested if inhibiting Unc-5 could reduce the increased frequency of spontaneous vesicle release when either Tomosyn or Complexin is also reduced. Consistent with prior studies in developing synapses, genetic depletion of Tomosyn or Complexin increased the frequency of spontaneous neurotransmission in adult motor synapses (Fig. 3a–f)(38). We next depleted Unc-5 together with simultaneous reduction of either Tomosyn or Complexin. Surprisingly, we found that depletion of Unc-5 suppressed the increased frequency of spontaneous vesicle release observed when either Tomosyn or Complexin are reduced (Fig. 3c,f). Conversely, Unc-5 depletion failed to suppress Tomosyn mediated potentiation of EJC amplitudes and quantal content (Fig. S5a–c). This suggested that, whereas Unc-5 dependant suppression of evoked amplitudes requires Tomosyn, Unc-5 mediated regulation of spontaneous neurotransmission acted ‘downstream’ of vesicular clamps. Given that vesicular release clamps interact directly with SNARE proteins which mediate vesicular membrane fusion, we speculated that synaptic Unc-5 might also interact directly with SNARE components at synapses(62, 63).

**Figure 3.**
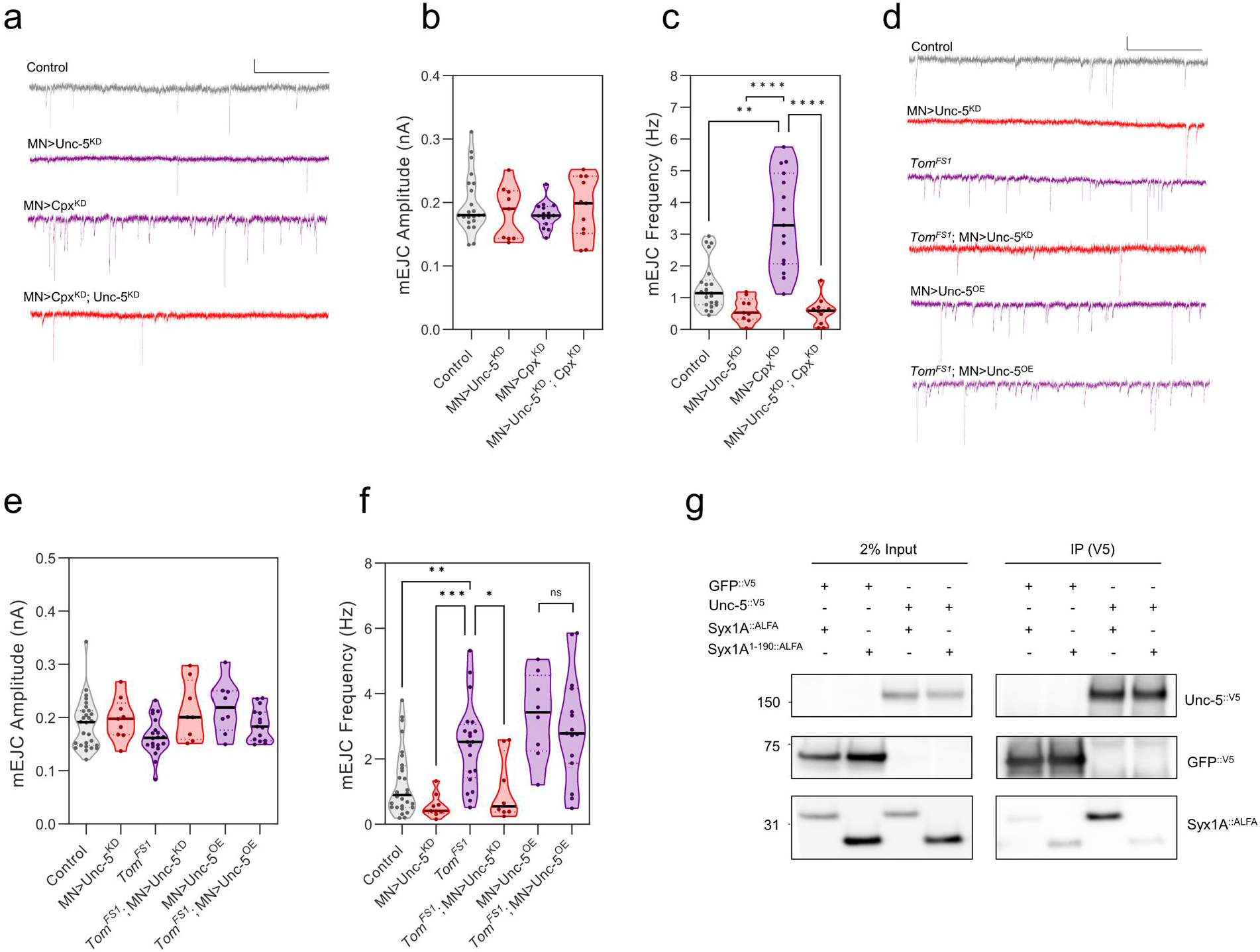
Unc-5 acts downstream of synaptic clamps and forms a complex with Syntaxin1A. **a**, Representative traces displaying excitatory glutamatergic mEJCs during conditional Unc-5 and Complexin depletion (Scale Bars: mEJC, 0.1 nA/1 s). **b**, Violin plots depicting no change in mEJC amplitude. **c**, However, Unc-5 depletion was able to suppress complexin mediated mEJC frequency increase (*n >* 11 recordings per genotype, Kruskal-Wallis test: *P <* 0.0001, Dunn’s multiple comparisons test: MN*>*CpxKD vs. MN*>*CpxKD; Unc-5KD, *P <* 0.0001). **d**, Representative traces displaying excitatory glutamatergic mEJCs during Unc-5 conditional depletion and Tomosyn deletion (Scale Bars: mEJC, 0.1 nA/1 s). **e**, Similarly to Complexin, no change in mEJC amplitude was detected. **f**, However, loss of function of Unc-5 was sufficient to deplete Tomosyn mediated mEJC frequency increase (*n >* 8 per genotype, Kruskal-Wallis test, *P <* 0.0001, Dunn’s multiple comparisons test: *Tom*FS1 vs. *Tom*FS1; MN*>*Unc-5KD, *P* = 0.013). Whereas, Tomosyn and Unc-5 gain of function were not additive. **g**, Unc-5 physically associates with Syntaxin1A via co-immunoprecipitation conducted from transgenic expression of Unc-5 and Syntaxin1A in *Drosophila* S2 cells. This interaction requires the Syntaxin1A C-terminal domain as evidenced by a dramatic decline of co-immunoprecipitation with truncated Syntaxin1A (Syx1A1-190). Violin Plots represent median & IQR. All detailed statistical analysis & genotype details are reported in Supplementary Table 1.

To test this hypothesis, we examined if Unc-5 protein could interact with the SNARE protein Syntaxin(64) in *Drosophila* S2R+ cells. We first confirmed previous results that Tomosyn directly interacts with Syntaxin by co-immunoprecipitation(65) (Fig. S5d). We then tested if Unc-5 also interacts with Syntaxin. We found that Syntaxin co-immunoprecipitated with Unc-5, confirming these proteins can form a complex (Fig. 3g). Tomosyn has previously been shown to interact with the C-terminal SNARE domain of Syntaxin(65) (*R. norvegicus*: AA191–266, *D. melanogaster* : AA195–257). We also confirmed this result, as deletion of this domain abolished co-immunoprecipitation of Tomosyn and Syntaxin (Fig. S5d). We then tested if Unc-5 also required this domain to interact with Syntaxin. We found, that similar to Tomosyn, Unc-5 also failed to co-immunoprecipitate with Syntaxin when its C-terminal domain was deleted (Fig. 3g). These data established that Unc-5 associates with the SNARE protein Syntaxin via the same region required for interaction with the vesicular clamp Tomosyn. This suggested the potential for a competitive interaction to regulate spontaneous neurotransmission. To test this, we examined the effects of increasing Unc-5 expression, which increases spontaneous neurotransmission, while simultaneously depleting Tomosyn, which also increases spontaneous event frequency when depleted alone. We found that increasing Unc-5 while also depleting Tomosyn failed to produce an additive increase in spontaneous neurotransmission frequency, as would be predicted if they functioned independently (Fig. 3f). Instead this result, consistent with our protein interaction data, suggests that Unc-5 and Tomosyn compete for association with Syntaxin to govern spontaneous neurotransmitter vesicle fusion.

### Unc-5 is essential for adult synaptic structural maintenance and behaviour

We have previously shown that declining levels of spontaneous vesicle release induces characteristic adult synaptic terminal bouton structural degeneration or fragmentation(38, 39). In this process larger synaptic bouton varicosities with multiple AZs progressively fragment into smaller units with fewer AZs iteratively until this subdivision results in boutons with only a single active zone. We have previously shown that the number of fully fragmented synaptic boutons increases steadily with age and is causally linked to a parallel age-dependent decline of the frequency of miniature neurotransmission events(38). As elevation of Unc-5 at adult synapses concomitantly increased spontaneous event frequency, we predicted that Unc-5 could also impact adult synaptic structural integrity. To examine this, we first assessed the effects of increasing synaptic Unc-5 on adult synaptic structures in old animals. When we compared animals expressing increased Unc-5 compared to age-matched controls, we found that the number of fragmented boutons was reduced by 43.31% (*P <* 0.01, Fig. 4a–b), consistent with increasing levels of Unc-5, and thus spontaneous release frequency, counteracting the synaptic structural consequences induced by the decline of miniature events with age(38, 39).

**Figure 4.**
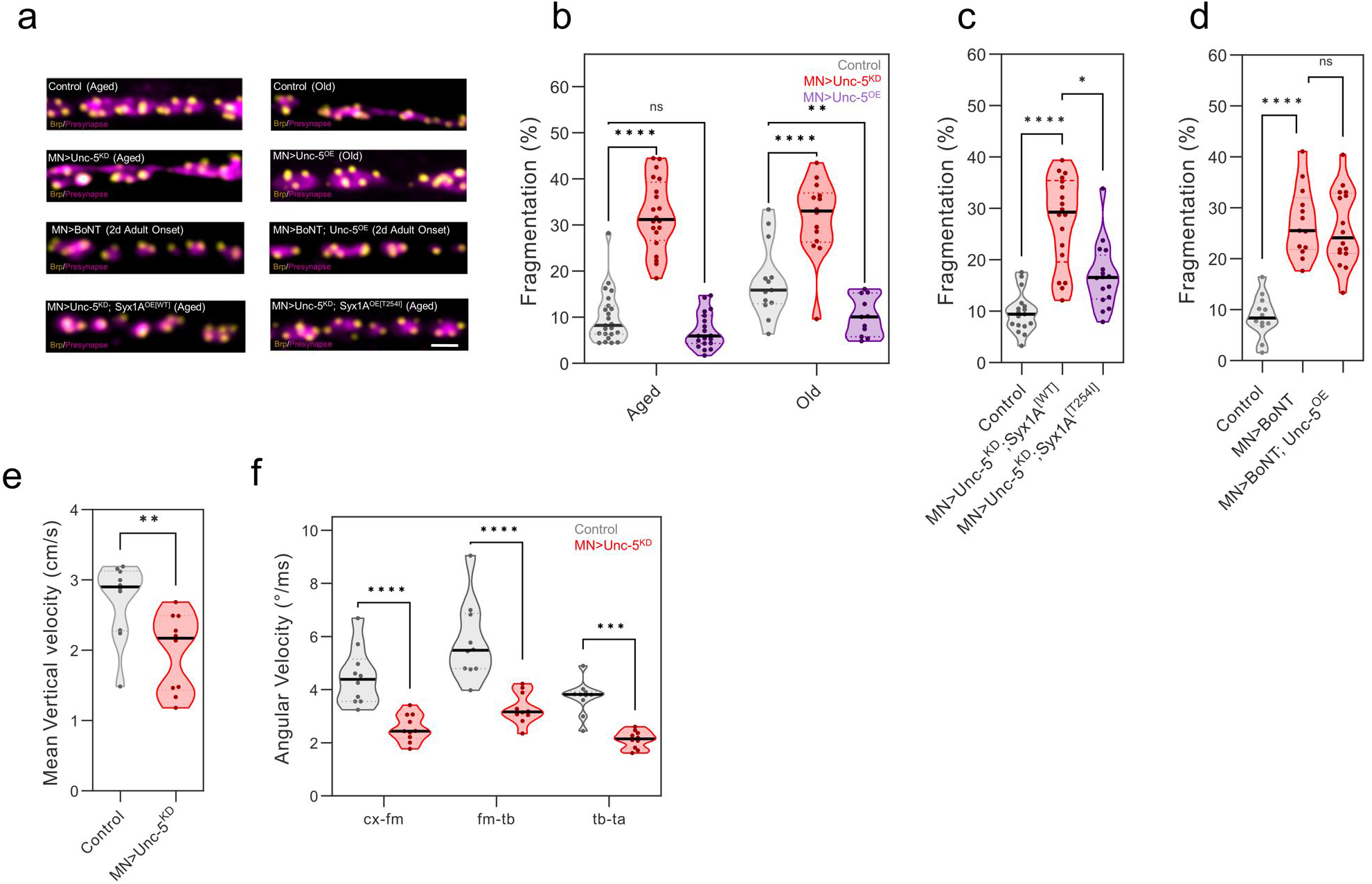
Synaptic Unc-5 depletion induces miniature dependent structural degeneration & behavioural deficits. **a**, Representative confocal images of Adult MVIM presynaptic boutons (magenta) and the AZ marker (brp) (Scale Bar: 3 µm). **b**, Violin plots showing that Unc-5 depletion increases the number of boutons containing a single AZ (fragmented) at middle age (20 days; pre-age-related structural decline) and old age (30 days; post-decline) (*n >* 11 NMJs per genotype, Dunnett’s multiple comparisons test, *P <* 0.0001). Whereas, increased Unc-5 levels can ameliorate age dependent fragmentation (*n >* 11 per genotype, Dunnett’s multiple comparisons test, *P <* 0.001, Control vs. MN*>*Unc-5OE). **c**, Unc-5 mediated structural depletion can be rescued through increased miniature neurotransmission via transgenic upregulation of Syx1AT254I (*n >* 16 NMJs per genotype, Dunn’s multiple comparisons test: MN*>*Unc-5KD; Syx1AWT vs. MN*>*Unc-5KD; Syx1AT254I, *P <* 0.05). **d**, Conversely, increased Unc-5 levels were unable to ameliorate structural degeneration induced through conditional BoNT expression (*n >* 13 NMJs per genotype, Dunnett’s multiple comparisons test, *P* = 0.86). **e**, Mean vertical velocity of climbing flies was reduced when Unc-5 was depleted (*n* = 100 animals, *N* = 10 vials per genotype, Student’s *t*-test, *P <* 0.01). **f**, This decline in locomotor performance was associated with a decline in limb angular acceleration at all leg joints analysed in walking animals. Identification of NMJ boutons was conducted at MVIM A2 using MN*<*UAS-mCherry and anti-VGlut for **b** and anti-VGlut alone for **c**,**d**. Violin Plots represent median & IQR. All detailed statistical analysis & genotype details are reported in Supplementary Table 1.

We next examined the effects upon synaptic structures of reducing synaptic Unc-5 levels in young animals. We found that adult onset depletion of Unc-5 caused a 222.41% increase in the number of fragmented synaptic boutons in these animals (*P <* 0.0001) (Fig. 4a–b). To exclude the possibility that these effects upon synaptic structures were mediated by activities of Unc-5 other than that upon miniature event frequency, we sought to increase the frequency of miniature events when Unc-5 levels were depleted. However, as we have shown above, depletion of synaptic vesicle clamps which normally induce increased spontaneous fusion events(44, 61), is suppressed by the reduction of Unc-5. Given this and building upon our data that Unc-5 interacts directly with Syntaxin, we generated a transgenic mutant form of Syntaxin (*Syx1A* ^*T* 254*I*^), based on a Syntaxin mutant allele that had previously been described to disproportionately increase spontaneous vesicle release(66, 67). We first confirmed that conditional overexpression alone of *Syx1A* ^*T* 254*I*^ neurons increased the frequency of spontaneous vesicle release (Fig. S6a–e). We then expressed Syx1A ^*T* 254*I*^ in adult while simultaneously depleting Unc-5. We found that the frequency of miniature events was increased by 506.03% (*P <* 0.001) when *Syx1A*^*T* 254*I*^ was co-expressed together with Unc-5 depletion relative to Unc-5 reduction alone (Fig. S6f–h). We then examined the structure of synaptic terminals in these two conditions. We found that the number of fragmented synapses when Unc-5 was depleted was reduced by 38.96% (*P <* 0.05) (Fig. 4c) when *Syx1A* ^*T* 254*I*^ was co-expressed vs. when Syx1A^*W T*^ was co-expressed. This result supported that the effects of Unc-5 reduction upon synaptic structures, 223 was mediated through a decline of miniature neurotransmission. While the frequency of miniature events decline with age as described above(38), they can also be (together with evoked release) depleted in young animals through the cleavage of Syntaxin via transgenic expression of Botulinum Neurotoxin(43). We expressed BoNT in adult neurons while simultaneously increasing expression of Unc-5. As predicted, increasing Unc-5 expression was unable to increase miniature event frequency in the presence of BoNT (Fig. S6k–l). When we examined the structure of synapses expressing both Unc-5 and BoNT, we found the levels of bouton fragmentation were not significantly different to animals expressing BoNT alone (Fig. 4d). Therefore, increasing or decreasing miniature event frequency by manipulating Syntaxin can modulate the effects of Unc-5 upon synaptic integrity establishing that these structural effects are mediated via altering spontaneous vesicle release.

As the reduction of Unc-5 in adult motor neurons had profound effects on synaptic structures as described above, we wondered if this degeneration also impacted adult motor ability and behaviour. We first measured motor ability using a negative geotaxis vertical climbing task when Unc-5 was depleted in all adult motor neurons versus controls. We found that the motor ability of Unc-5 depleted animals was significantly impaired compared to controls with a 27.17% (*P <* 0.01) decrease in climbing ability (Fig. 4e). To understand the cause of this deterioration in more depth, we used deep learning(68, 69) to evaluate the individual limb kinematics of Unc-5 depleted animals and controls (Fig. 4f). We found that the mean limb angular velocity of Unc-5 depleted animals was dramatically reduced (*−* 54.58% pooled average, *P <* 0.001) compared to controls, suggesting an inability to perform and sustain intensive motor behaviours. In sum, Unc-5 is required in adult synapses not only to sustain miniature events, but as a consequence also preserve adult synaptic structural integrity and behavioural capacity.

## Discussion

Our results establish that the release of individual synaptic vesicles is a regulated process and not a spontaneous epiphenomenon as previously thought. We show that the synaptic levels of the transmembrane signalling protein Unc-5 bidirectionally governs the frequency of these events. We find that Unc-5 is enriched at AZs that release synaptic vesicles in the absence of action potentials and that this localization is dependent upon synaptic cleft HSPG activity. We further show that Unc-5 protein associates with the SNARE Syntaxin via the same region necessary for interaction with the synaptic clamp protein Tomosyn, supporting a mechanism in which the association of Unc-5 with Syntaxin competes with and prevents the clamping activity of Tomosyn to promote vesicle release. Consistent with this, in our experiments, only the direct manipulation of Syntaxin, via point mutations or BoNT toxin disruption, could alter the effects of Unc-5 upon spontaneous neurotransmission. In contrast, the parallel alteration of evoked neurotransmission observed with Unc-5 depletion could be modified by changing Ca^2+^ or Tomosyn levels, indicating that the effects of Unc-5 on evoked neurotransmission is subject to independent regulation. Consistent with this, evoked neurotransmission remains unchanged when Unc-5 was overproduced. Together, these findings identify Unc-5 as an AZ segregated synaptic regulator that selectively promotes spontaneous vesicle release through interaction with Syntaxin.

Our results demonstrate that the release properties of adult *Drosophila* synaptic AZs are heterogeneous, consistent with prior studies in developing *Drosophila* synapses and with recent studies in other systems(25–31). Levels of Unc-5 appear critical for this heterogeneity of release probability, at least at release sites that produce miniature events, suggesting synaptic release sites should be reconsidered to be functionally but also molecularly diverse, and not viewed as uniform and interchangeable. Our data adds further evidence that ‘spontaneous’ release events are, like evoked release, an essential component of neurotransmission that is critical for brain function, at a minimum for the structural maintenance of synapses. Given our evidence that these fusion events are regulated, we suggest miniature neurotransmission (rather than spontaneous) to be the more accurate term to describe this release modality. When miniature neurotransmission is depleted, as we show here when synaptic Unc-5 is reduced, synaptic terminal structures degenerate and behavioural capacity is compromised. This is consistent with other studies, including by us, that have revealed unique roles for miniature events in vivo that are not shared with evoked neurotransmission(34–38). Indeed, the independence of miniature neurotransmission from evoked release may make these miniature events uniquely suited for longer-term synaptic signalling roles, such as in synaptic structural maintenance, which can be uncoupled from the evoked firing activity of the neuron. Beyond identifying Unc-5 as a distinct regulator of miniature neurotransmission, our spatial imaging reveals that nearly one quarter of all synaptic active zones are exclusively tasked with generating miniature events, persuasive evidence for their fundamental importance to neuronal function in vivo.

## Acknowledgments

This work was supported by the Swiss National Science Foundation grant numbers: 31003A_179587 and 320030-232324 to B.M.

## Author Contributions

S.V. & B.M. conceived and designed the study. S.V. designed, implemented and analysed the electrophysiology, live imaging, IHC, conducted climbing behavioural experiments and aided in limb velocity analysis. E.R. generated transgenic constructs and scientific illustrations. C.P. aided construct generation and carried out cell transfection and co-immunoprecipitations. R.S. implemented and analysed limb velocity data. M.V.C. conducted WBs and analysed climbing behavioural data. S.V. and B.M. wrote the manuscript. All authors reviewed and approved the final manuscript.

## Materials and Methods

### *Drosophila* stocks & husbandry

*GAL4 Lines:* OK6-GAL4(70), VGMN-GAL4(71). *UAS-RNAi lines:* UAS-Unc-5*RNAi* (BDSC#33756), UAS-luciferase^*RNAi*^ (BDSC#31603), UAS-Cpx ^*RNAi*^ (BDSC#42017). *UAS-Upregulation lines:* UAS-Unc-5^::*HA*^ (Gift from Greg Bashaw), UAS-degradFP^5*R*^ (this study), 20xUAS-6xmCherry-HA(72), UAS-BoNT-C(43). *Mutant Lines:* SynapGCaMP6f(27), *Unc-5* 5^::*smV* 5^ (this study), *Brp*::*RFP* (this study), *Unc-5*::*GFP* (BDSC#60547), tub-GAL80*ts* (73), *TomFS*1 (44), netABΔ(54), 8 ^*GT*1^ *unc-5* (74), *sflGFP* (BDSC#12782)(59), *sfl* (BDSC#63165)(75), *sflDf* (Df(3L)ED211, BDSC#8063).

*Drosophila* stocks were reared under standard rearing conditions. Temporal control of targeted expression of transgenes was conducted as previously described(38). Briefly, male flies were raised at restrictive temperature (18°C) and switched to permissive temperature (29°C) after at least 5 days. Due to the time course of RNAi mediated knockdown, experiments were conducted on flies considered aged (20 days) or old (30 days) as indicated.

### Transgenic Reagents

#### Brp::RFP

To obtain a conditionally tagged Brp protein with a red fluorescent protein, a blownOUT stop cassette(76) and a mScarlet tag(41) were inserted in genomic DNA at the C-terminal of Brp-RD variant, using CRISPR/Cas9 based genome editing(49). To facilitate the screening of transformed flies, a DsRed cassette was added after the stop codon of mScarlet coding sequence. All PCR amplifications were performed using Platinum Superfi polymerase (Invitrogen) and assembled using HIFI technology (New England Biolabs). First, 5 fragments were assembled as follows: (1) 1198 bp Homology Arm 1 (HA1) flanking the region upstream of the 5’ gRNA target site and (2) the region between 5’ gRNA target site and the stop codon of Brp gene were amplified from *Drosophila* genomic DNA. A modified PAM sequence was inserted on overlapping primers. (3) The DsRed selection cassette, 3xP3-Hsp70pro-dsRed2-SV40poly, flanked by two inverted terminal repeat sites, was PCR amplified from pHD-sfGFP Scarless dsRed (Addgene #80811). KpnI and XmaI restriction sites were added on the forward primer for later insertion of the stop cassette and the mScarlet tag. (4) The region just after the stop codon of Brp gene until the 3’ gRNA target site and (5) the 1079 bp Homology Arm 2 (HA2) flanking the region downstream the 3’ gRNA target site, were amplified from *Drosophila* genomic DNA. The assembly was topo cloned in zero-blunt end pCR4 vector (Invitrogen) giving rise to pCR4 Brp-KpnI-XmaI-DsRed. Secondly, the BlownOUT stop cassette flanked by 2 modified Rsrt sites was PCR amplified from pJFRC168 rstr-2xstop-rstr*>*Myr::RFP (Addgene #32143) and inserted by digestion-ligation using KpnI and XmaI in pCR4 Brp-KpnI-XmaI-DsRed, giving rise to pCR4 Brp-BlownOUT Stop-XmaI-DsRed. Finally, the Scarlet sequence was PCR amplified from pF3BGX-mScarlet-I (Addgene #138391) and inserted by digestion-ligation using XmaI restriction site in Brp-BlownOUT Stop-XmaI-DsRed, giving rise to the final construct pCR4 Brp-BlownOUT Stop-Scarlet-DsRed. Two gRNA (5’ and 3’) sequences targeting each side of the knock-in insertion (5’gRNA 5’-GGTTGTGTTTTCAGCGAT-3’, 3’gRNA 5’-GGGTCTTACAGATGGGAA-3’) were selected using the FlyCRISPR algorithm (http://flycrispr.molbio.wisc.edu/) and were predicted to have minimal off-targets. Both gRNA sequences were individually inserted into separate pCFD3 vectors (Addgene #49410) using the KLD enzyme mix (New England Biolabs). To generate the transgenic flies the HDR template plasmid and the two gRNA plasmids were sent for microinjection into [nos-Cas9]attp2 (BDSC#54591) embryos at Genetivision, USA.

#### *Unc-5*^::*smV* 5^

Using CRISPR/Cas9 based genome editing(49), a Spaghetti-monster-V5 (smV5) tag was inserted in the Unc-5 gene, 29 amino acids downstream of the start codon. An excisable DsRed selection cassette was inserted in an intron, 173 bp upstream of Unc-5 5’UTR. All PCR amplifications were performed using Platinum Superfi polymerase (Invitrogen) and assembled using HIFI technology (New England Biolabs). The template construct was generated in three steps. First, the three following segments were assembled together; (1) 1033 bp HA1 flanking the 5’ region upstream of Unc-5 gene until 5’ gRNA target site, (2) the region between 5’ gRNA target site and 3’ gRNA target site and (3) 1103 bp HA2 flanking the region downstream of 3’ gRNA target site were amplified from *Drosophila* genomic DNA and cloned in zero-blunt end pCR4 topo vector (Invitrogen) generating the construct pCR4-HA1-mutated PAMs-HA2. Modified PAM sequences were inserted on overlapping primers. Second, the three following segments were assembled: (4) 3xP3-Hsp70pro-dsRed2-SV40polyA selection cassette, flanked by two PiggyBac sites, was amplified from pHD-sfGFP Scareless dsRed (Addgene #80811), (5) the region between the DsRed insertion site and the tag insertion site was amplified from *Drosophila* nos-cas9 (attp2) genomic DNA and (6) the smV5 tag was amplified from pCAG smFP V5 (Addgene #59758). The full assembly was then gel purified (GeneJET, Thermo Fisher) generating an insert containing the DsRed cassette and the tag. Finally, the vector pCR4-HA1-mutated PAMs-HA2 was PCR amplified from the DsRed cassette insertion site through the whole vector backbone until the tag insertion site. The vector and the insert (DsRed cassette – tag) were assembled by HIFI, generating the final construct pCR4 DsRed-smV5-Unc5. Two gRNA sequences targeting each side of the knock-in insertion (5’gRNA 5’-GGGTCGTCGGACGGGGTCGT-3’, 3’gRNA 5’-GGATGCTATTGACCCACTCA-3’) were selected using the FlyCRISPR algorithm (http://flycrispr.molbio.wisc.edu/) and were predicted to have minimal off-targets. Both gRNA sequences were individually inserted into separate pCFD3 vectors (Addgene #49410) using the KLD enzyme mix (New England Biolabs). To generate the transgenic flies, the HDR template plasmid and the two gRNA plasmids were sent for DNA microinjection into [nos-Cas9] attp2 (BDSC 54591) embryos at Genetivision, USA.

#### UAS-Syx1AWT & UAS-Syx1AT 254I

Constructs were synthesised with the addition of the Syn21 5*′*UTR sequence to enhance expression and inserted into pBID2_UASC(71) (Genscript, USA). Embryo injection and construct integration was targeted to landing site VK05 for both constructs (Genetivision, USA). The precise Syx1A modifications are described in Supplementary Table 3.

#### *UAS-degradFP*^[5*R*]^

To degrade GFP tagged proteins, a nanobody against GFP was fused to the *Drosophila* F-box domain, *slmb*(77).

In order to reduce degradation of the peptide by the proteasome machinery, we converted 5 Lysines into Arginine (K3R, K81R, K122R, K162R, K182R) and codon optimised this sequence for *Drosophila*. In addition to these mutations, we added a Syn21 translational enhancer sequence directly upstream of the start codon and a V5 tag on 5’ end. This sequence was synthesised by IDT, Switzerland.

The GFP nanobody was amplified by PCR from pCS2 mAID_vhhGFP(77) (Addgene #117718). V5-*slmb* and GFP Nb fragments were assembled using Hifi technology generating degradFP^[*R*5]^. The assembled sequence was cloned into pCR8GW-TOPO vector and then transferred in pBID2-UAS (71) vectors using LR II clonase (Invitrogen), resulting in pBID2-20xUAS-V5-degradFP^[5*R*]^. To generate the transgenic flies the plasmids were sent for DNA microinjection into embryos using the PhiC31 integration at Genetivision, USA. The 20xUAS-V5-degradFP^[5*R*]^ construct was inserted at the genomic position 66B4, chromosome 3 (fly strain JK66B).

#### *S2 expression constructs Syx1A*::*ALFA, Syx1A*1*−*190::*ALFA, Tom*::*M yc, Unc-5*::*V* 5

A Syn21 translational enhancer sequence was incorporated via the forward primer and positioned immediately upstream of the start codon in all constructs. Each construct coding sequence was amplified by PCR from the templates indicated: (1) *Syx1A-ALFA*: Syntaxin1A cDNA was amplified from DGC Gold collection(78). An ALFA tag was introduced before the stop codon using the reverse primer. (2) The truncated Syx1A(1–190) construct was generated by PCR amplification of the corresponding coding region from the full-length Syx1A-ALFA plasmid and cloned using standard molecular cloning techniques. (3) *Tom13A-6xMyc*: Tomosyn 13A-6xMyc cDNA was directly amplified from genomic DNA of Tomosyn 13A-6xMyc line(44). (4) *smV5-Unc-5*: Unc-5 cDNA and the Ruby_smFP_V5 tag were amplified from the genomic DNA of our CRISPR line smV5-Unc5. Both fragments were assembled using Hifi technology. All tagged coding sequences were first cloned into pCR8GW-TOPO vector and then transferred in the modified S2 cell expression vector, pExpreS2.1-Gateway(79), using LR II clonase (Invitrogen) resulting in pExpreS2.1 Act-Syx1A-ALFA, pExpreS2.1 Act-Tom13A-6Myc and pExpreS2.1 Act-smV5-Unc-5 constructs. All constructs were verified by sequencing (Microsynth AG, Switzerland).

### Cell line husbandry & co-immunoprecipitation

*D. melanogaster* S2R+ (FBtc0000150) were obtained from the DGRC and cultured in Schneider’s *Drosophila* medium (Bioconcept) supplemented with 10% FBS (Sigma) and 1% Penicillin-Streptomycin (Sigma). Effectene (Qiagen) was used for plasmid transfection according to the manufacturer’s protocol. 48 hrs after transfection cells were collected, washed with PBS and pelleted by centrifugation at 400 g for 10 min. The cell pellet was resuspended in 1 ml of lysis buffer (50 mM Tris-HCl at pH 7.4, 150 mM NaCl, 0.5% NP-40) supplemented with protease and phosphatase inhibitors and rotated head over tail for 30 min at 4°C. Protein lysates were centrifuged at 18,000 g for 10 min at 4°C to remove cell debris and protein concentration was determined using the Micro BCA Protein Assay Kit (ThermoFisher) following manufacturer’s protocol. For immunoprecipitation, 2 mg of protein lysate was incubated with 25 µl of Pierce Protein G magnetic beads (Thermo Scientific) coupled with 2 µg of mouse anti-V5 (1:1000, Invitrogen R960-25) or mouse anti-Myc (1:10000, Millipore 05-419) in 900 µl of lysis buffer and rotated head over tail overnight at 4°C. The beads were washed three times with 900 µl of lysis buffer. Immunoprecipitated complexes were eluted by incubation in 1*×* SDS loading buffer for 10 min at 70°C and analysed by Western Blot using mouse anti-V5 (1:1000, Invitrogen R960-25), mouse anti-Myc (1:10000, Millipore 05-419) and camelid HRP-conjugated anti-ALPHA (1:2000, Abnova RAB00932).

### Western Blots

*Drosophila* heads were lysed using RIPA buffer supplemented with cOmplete™ ULTRA Tablets (Roche) and PhosSTOP™ (Roche) to prevent protein degradation and dephosphorylation. Protein concentration was determined using the Micro BCA Protein Assay Kit (ThermoFisher) following manufacturer’s protocol. Equal amounts of protein lysates were mixed with 4 *×* loading buffer and heated at 95°C for 5 minutes before electrophoresis. Samples were separated on Novex™ WedgeWell™ 4–12% Tris-Glycine Gels (Invitrogen) using MES SDS Running Buffer 20 *×* (Life Technologies) and transferred onto PVDF membranes. After blocking in 5% non-fat milk in PBST (PBS + Triton 100X 0.1%) for 1 hour at RT, membranes were incubated overnight at 4°C in blocking solution with primary antibodies targeting the proteins of interest (mouse anti-V5 Invitrogen P/N46-0705 1:1000; mouse anti-tubulin Sigma T6199 1:10,000). Membranes were washed with PBST 3 *×* 15 minutes before incubation with secondary antibodies for 2 hours at RT (anti-mouse HRP Jackson Immuno Research AB_10015289 1:10,000). Membranes were washed with PBST 3 *×* 15 minutes before revelation. Protein bands were detected using enhanced chemiluminescence with the WesternBright™ Sirius WB detection kit (Advansta) and imaged using the Amersham Imager 680.

### Morphology & live imaging

#### Primary antibodies

chicken or rabbit anti-GFP (1:1000; AbCam), rabbit anti-RFP (1:500; Clonetech), mouse anti-Brp (Nc82, 1:100; DSHB), chicken anti-VGlut(38) (1:100), mouse anti-tubulin (1:100, DSHB), chicken anti-V5 (1:500, Invitrogen), mouse anti-V5 (1:500, Novus Biologicals), guinea pig anti-GluRIID(38) (1:500). *Secondary antibodies:* goat anti-chicken (Alexa-488, 555 or 647; 1:400, Life Technologies), goat anti-mouse (Alexa-Cy3 or 647; 1:400, Life Technologies), goat anti-rabbit (Alexa-488, 546 or 647; 1:400, Life Technologies), goat anti-guinea pig (647, 1:400, Life Technologies), Alexa Fluor 594-HRP (1:400; Jackson ImmunoResearch).

#### Adult and embryonic sample preparation

Adult MVIM terminal neuromuscular junction dissection and immunohistochemistry was conducted as previously described(38). Samples were mounted in ProLong Diamond antifade mountant (Invitrogen). Embryos were collected and staged at 25°C on apple agar plates supplemented with yeast paste. Standard methods were used for dechorionation, removal of the vitelline membrane and fixation(80). Embryos were stored in 100% ethanol at *−*20°C before IHC labelling and were mounted in VectaShield (Vector Laboratories) prior to imaging.

#### Fixed tissue image acquisition & analysis

Fixed tissue adult motor synapses and embryos were acquired on a Nikon Yokogawa Spinning Disk Confocal CSU-W1 with a Nikon CFI Plan Fluor 40 *×* Oil, 60 *×* water immersion or 10 *×* water immersion objective and denoised using inbuilt denoising software prior to manual analysis (Nikon, Japan). Fragmentation analysis was quantified as previously described(38). For automated colocalization analysis, images were deconvolved with Huygens deconvolution software (SVI, Netherlands) prior to analysis. Colocalization between Unc-5 (Channel 1) and the AZ marker (Brp, Channel 2) was quantified using Manders’ Overlap Coefficient M2, representing the colocalization of Brp signal with the Unc-5 signal. Measurements were calculated using the Fiji plugin, JACoP (EPFL, BIOP) computed from quantification of mean M2 values from 5–10 ROIs taken from single Z-planes of AZs for each NMJ. Manual threshold values for Unc-5 and Brp channels were determined from independently stained single-channel control images.

#### Live imaging and analysis

Live imaging of SynapGCaMP^6*f*^ mediated events was performed on a Nikon Yokogawa Spinning Disk Confocal CSU-W1. Adults were dissected in AHL saline (0 mM Ca^2+^ ) to expose the ventral midline A2 MVIM NMJs and image acquisition was recorded in 1.5 mM Ca^2+^ AHL saline. We restricted live imaging recordings to a maximum of 1 minute per recording to minimize photobleaching. Post-acquisition Z-stack projections of Brp^::*RFP*^ and GCaMP channels were conducted for registration and structural reference. Spontaneous and evoked neurotransmission were recorded sequentially from the same NMJ. Spontaneous activity was recorded for 1 minute, evoked events were simulated for 30 s (1 ms/0.5 Hz) through direct nerve stimulation.

Prior to analysis images were denoised using Nikon AI denoising software and background subtracted to assist manual event identification. Brp^::*RFP*^ puncta were labelled in Z-stack acquisitions and AZ ROIs were drawn based on their positions within corresponding live GCaMP recordings of the same NMJ. GCaMP events were identified by visualising fluorescence changes (Δ*F* ) in z-axis intensity profiles at individual AZ ROIs. Evoked activity was confirmed by the presence of rapid muscle twitches; analysis was restricted to time-locked stimulation periods. NMJs showing reduced evoked responses, below safety threshold and lacking muscle twitches, were validated by post hoc high frequency stimulation (100 Hz) to confirm nerve innervation.

AZs were classified as follows: Somnolent AZ (0:0, spontaneous:evoked), spontaneous AZ ( *≥* 1:0), evoked AZ (*≥* 0: 1), mixed AZ ( *≥* 1: *≥* 1). Spontaneous release probability was expressed as Hz and evoked release probability as response per stimulation. Covariable somnolent AZ populations were excluded from release probability analysis. Release probability values were averaged amongst recorded AZs per recording.

### Electrophysiology

*dSEVC recordings* were conducted as previously described(38). Briefly, recordings were conducted in 1.5 mM Ca^2+^ AHL saline unless otherwise stated. Voltage clamp recordings of adult neuromuscular glutamatergic synaptic currents were carried out using borosilicate glass electrodes (1B120F-4, World Precision Instruments, Florida, USA). Recording electrodes were pulled (Sutter P-1000, California, USA) to resistances of between 20–25 M Ω and filled with 3 M KCl. Suction electrodes (GC120T-10, Harvard Apparatus, Kent, UK) were fire polished to minimize damage to the nerve to a diameter of *∼* 2–3 µm and filled with extracellular saline. EJCs were evoked (1 ms/0.5 Hz or 100 Hz for HFS recordings) with a 2100 isolated pulse stimulator (AM Systems, Washington, USA). mEJC and EJC recordings were made from individual muscle fibres of the A2 MVIM abdominal muscle segments. Recordings were made using an Axoclamp 900A amplifier (Molecular Devices, Sunnyvale, CA). Muscle fibres were held at *−*60 mV, recordings sampled at 20 kHz and lowpass filtered at 0.5 kHz using pClamp 11 (Molecular Devices, Sunnyvale, CA). Recordings were only accepted for analysis if muscle input resistance was *>*10 M Ω and injected current was *<*2.5 nA. mEJC amplitude and frequency were quantified from 2 minute recordings using automated threshold detection(81). Only events *>*100 pA were included for analysis corresponding with *>*7.5 *×* average recording noise to mitigate false positive event detection. EJC amplitudes were averaged from 10 EJCs per recording. Combined live imaging and electrophysiology was conducted as above and recorded simultaneously.

### Behaviour

*Negative geotaxis motor ability assay:* Male flies were maintained in groups of 10 per food vial and transferred to fresh food every 2 days throughout the experiment. Flies were transferred to empty transparent vials (15 cm height, 2.5 cm diameter) in groups of 10 and allowed to acclimatize for 30 minutes at room temperature. To initiate negative geotaxis, a custom built holder was lifted to a fixed height and released three times in succession. Climbing behaviour was recorded using a Pi-Camera connected to a Raspberry Pi and operated via the open-source PiCameraApp.py interface. Videos were captured at 30 frames per second for at least 25 seconds per trial under uniform lighting conditions. Mean vertical climbing velocity was calculated per group of 10 flies during the first 2 seconds of climbing using automated FreeClimber software(82). Data are presented as mean vertical velocity (cm/s).

#### Limb Kinematics

Flies were tethered to a holder with UV light hardened dental glue and placed into the behavioural arena. We recorded the flies from a side view camera (Basler, acA1920-155um, Germany) of 94 mm *×* 94 mm focal length with a 1.00 *×* InfiniStix lens (Proximity Series, 94 mm/1.00 *×*, USA) that was synchronized using an Arduino board hardware trigger. Animals were illuminated with infrared (850 nm) ring lights (CSS, LDR2-74IR2-850-LA, Japan). We recorded the flies behaving on the treadmill at 100 frames per second for 1 minute.

We generated 2D pose estimations of the fly legs from our recordings using DeepLabCut(68). In our training set, we labelled 17 key points: the head, abdomen tip, as well as 5 points per leg for the front, middle and back legs on the left side of the fly: on the joints of the body-coxa, coxa-femur, femur-tibia, tibia-tarsus, and tarsus tip. Our training dataset included 95 flies of different genotypes with 40 frames per fly labelled to ensure *>*99% accuracy for every fly. For training, we used an effnetB6 network, a training fraction of 0.95, a filtered p-cut-off of 0.6, and we trained the network for 40,000 iterations. We then processed the tracked joint coordinates over time using the dlc2kinematics (https://github.com/AdaptiveMotorControlLab/DLC2Kinematics) python library to compute angular velocity for each fly during the trials. We filtered the data to include only when the fly was moving by using a threshold of an average absolute angular velocity of 1 across legs. From this movement dataset, we computed average angular velocity per joint across genotypes for comparison.

### Statistical analysis and reproducibility

Statistical analyses were performed in Prism v.10.6.1 (GraphPad). Normality was assessed using the Shapiro-Wilk test and equality of variance with the Brown-Forsythe test. If either assumption was violated, non-parametric alternatives were used. Otherwise, statistical significance was assessed by two-tailed *t*-test, one-way or two-way ANOVA. ANOVA based post-hoc Dunnett’s test was conducted when multiple comparisons were required. All sample sizes are indicated in figure legends. Sample sizes are denoted by ‘*n*’ in text, figure legends and tables representing mean values per single NMJ for morphological analysis, single recording from live imaging or a single muscle fibre for electrophysiology; a maximum of two NMJs, recordings or muscle fibres were taken from biologically independent samples during data acquisition.

## Ethics statement

This study employed *Drosophila melanogaster* which is not subject to institutional ethical approval. All relevant regulations for the use of genetically modified *Drosophila* were complied with.

## Data availability

Source data is available upon request. All other reagents including *Drosophila* strains are available upon request.

---

**Figure S1.**
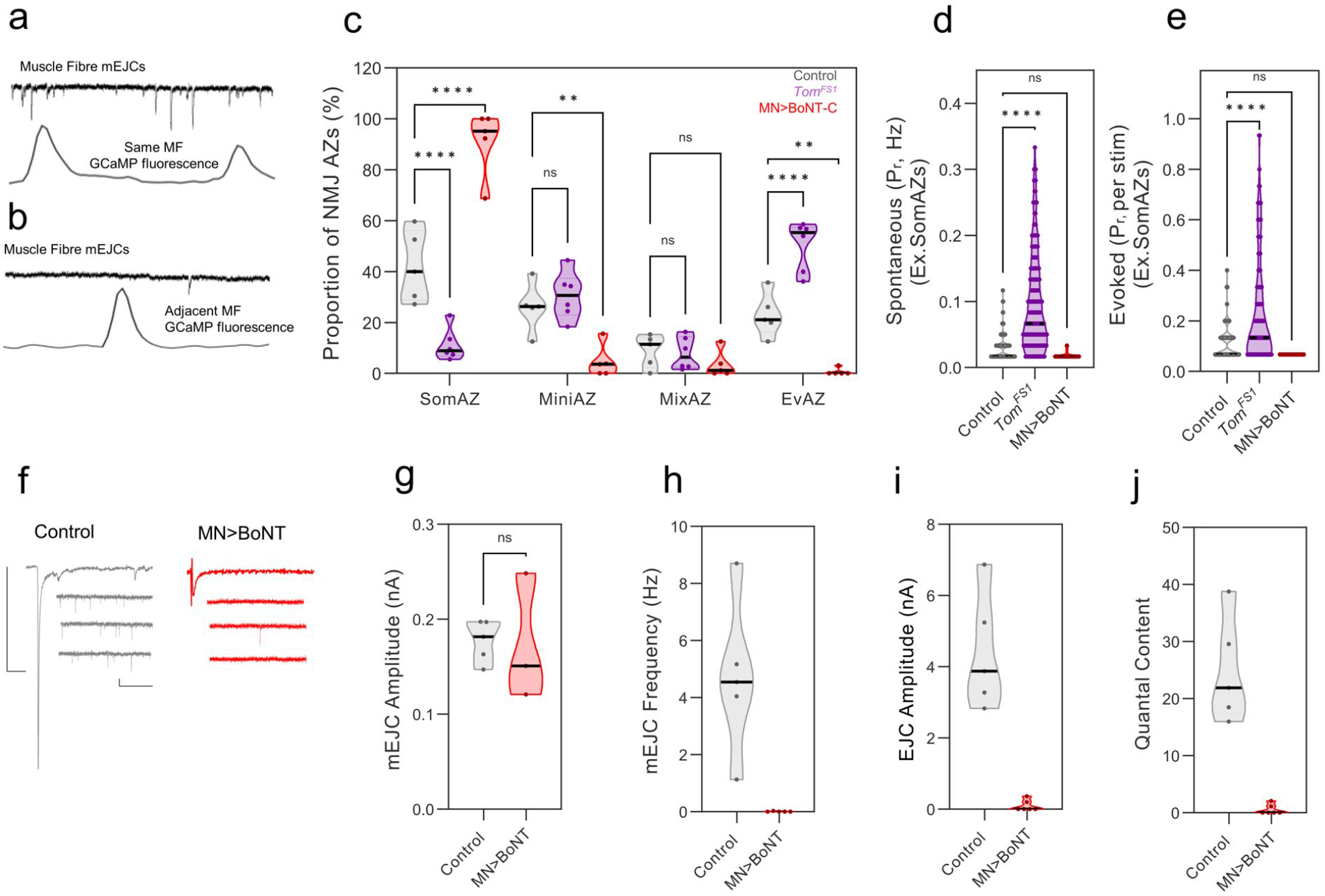
Validation of live neurotransmission spatial imaging tools. **a**, Representative traces of simultaneously acquired mEJC measurements representing total synaptic input into the recorded muscle fibre together with GCaMP fluorescence signals observed at a single AZ. Plots show observed amplitude correlate within the same muscle fibre and **b**, showing imaged adjacent muscle fibre displaying no relationship with the acquired GCaMP signal. **c**, Violin plots representing strong bi-directional alteration of observed AZ functional identity when neurotransmission is genetically challenged to increase spontaneous and evoked release using the Tomosyn deletion (*Tom*^FS1^) or decrease spontaneous and evoked release through conditional Botulinum toxin expression (MN*>*BoNT) (*n >* 5 recordings per genotype, Two-Way ANOVA, *P <* 0.0001). **d**, Violin plots showing altered spontaneous and **e**, evoked *Pr* rate change at individual AZs. These modifications to function were validated using electrophysiology both *Tom*^FS1^ mutants (**Fig. 3d–f & Fig. S5a–c**) and BoNT (Scale Bars: EJCs, 2 nA/30 ms mEJCs, 0.1 nA/500 ms) **f–j**. Violin Plots represent median & IQR. All detailed statistical analysis & genotype details are reported in Supplementary Table 2.

**Figure S2.**
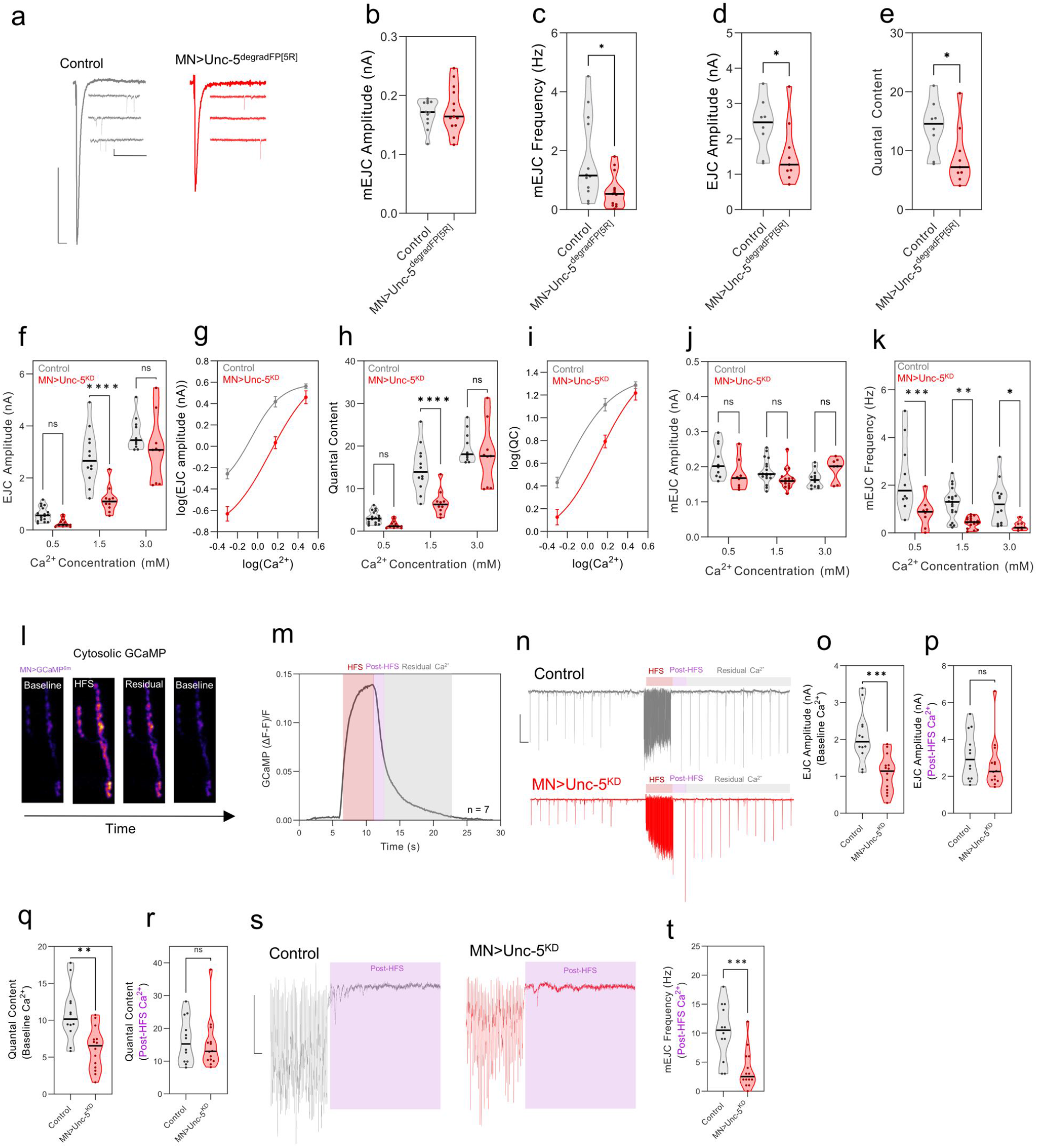
Unc-5 modulates evoked release independently of its effect on spontaneous release. **a–e**, Protein targeted nanobody mediated knockdown of Unc-5 (MN*>*Unc-5^degradFP[5R]^) recapitulates RNAi mediated functional phenotypes (Scale Bars: EJCs, 1 nA/30 ms mEJCs, 0.1 nA/500 ms). **f–i**, Violin plots and minimal Ca^2+^-dependence curves (three points: 0.5, 1.5, and 3 mM extracellular Ca^2+^) showing EJC amplitude and quantal content. Analysis is sufficient to show a right shifted curve showing partial saturation of evoked response at elevated Ca^2+^ to a similar level as control in Unc-5 depletion. **j–k**, Whereas, Unc-5 mediated mEJC frequency decline is independent of extracellular Ca^2+^. **l**, Representative GCaMP images of synaptic boutons at the MVIM A2 terminal expressing cytoplasmic GCaMP6m during 5 s/100 Hz stimulation in 1.5 mM Ca^2+^ extracellular saline. **m**, Quantification of GCaMP fluorescence dynamics with overlays highlighting HFS period (red), an initial elevated post-HFS residual Ca^2+^ period (magenta) followed by a longer decay of residual Ca^2+^ (grey). **n**, Representative traces showing EJC response to HFS (stimulation paradigm: 0.5 Hz – 100 Hz/5 s – 0.5 Hz, Scale Bar: 1 nA/2 s). **o–r**, When Unc-5 levels are depleted, initial EJC amplitude and QC is lower than controls (*n >* 12 recordings per genotype, Mann Whitney test, EJC, *P <* 0.001, QC, *P <* 0.01). However, EJC amplitude is rescued to 85.34% of control values (93.33% QC rescue) when quantified in the post-HFS residual calcium period, where cytoplasmic Ca^2+^ is elevated (*n >* 12 recordings per genotype, Mann Whitney test, *P* = 0.7). **s**, Representative traces displaying HFS induced mEJCs released immediately proceeding HFS (Scale Bar: 1 nA/50 ms). **t**, Despite EJC and QC rescue observed the same recordings, the rate of mEJC release was maintained as dramatically reduced relative to controls when Unc-5 was depleted (*n >* 12 recordings per genotype, Mann Whitney test, *P <* 0.001). Violin Plots represent median & IQR. All detailed statistical analysis & genotype details are reported in Supplementary Table 2.

**Figure S3.**
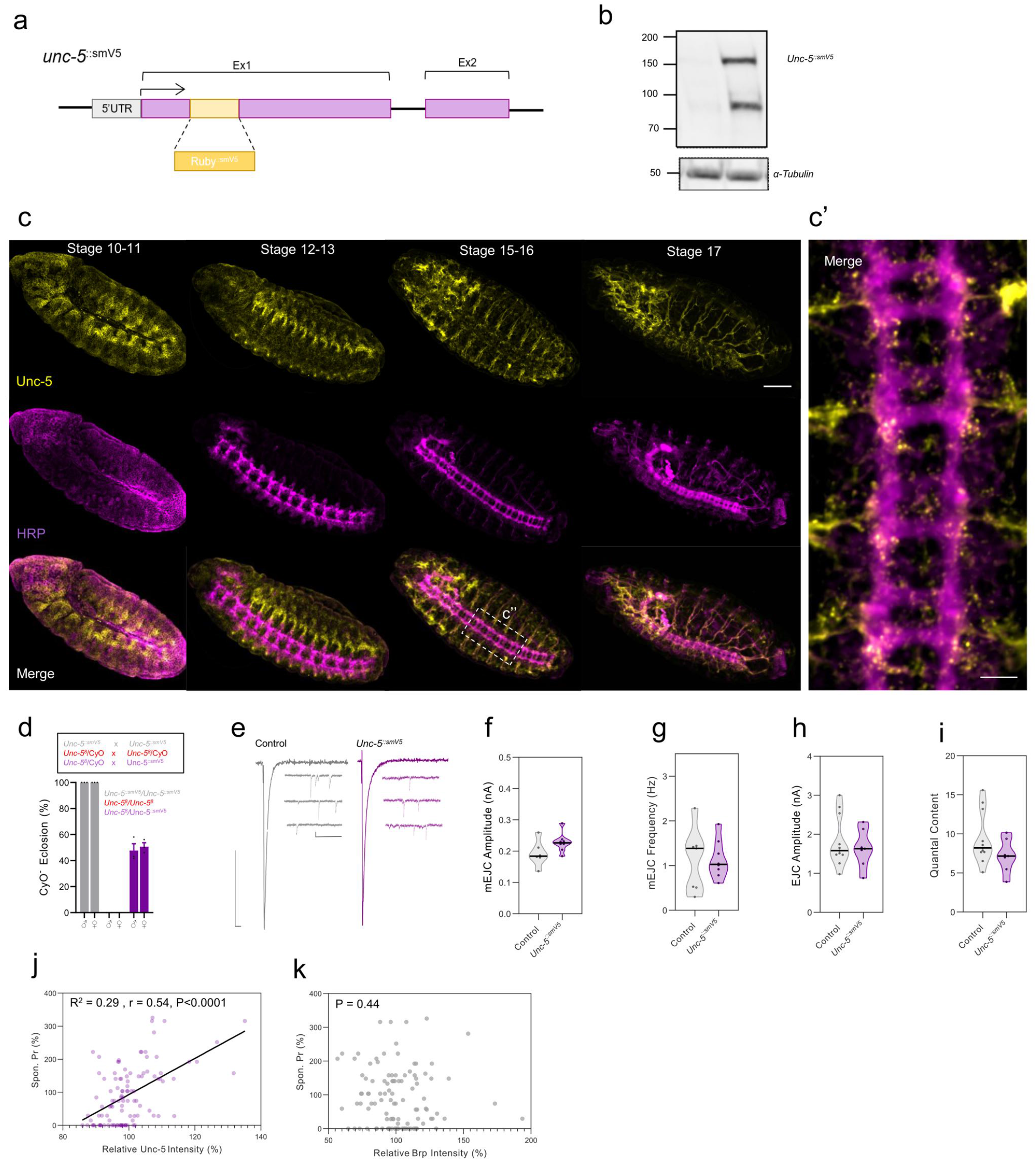
Validation of genome edited epitope tagged Unc-5::smV5. **a**, Schematic representing location of the Unc-5^::smV5^ tag within the first exon of Unc-5. **b**, Western blot showing predicted weight for Unc-5^::smV5^ (Unc-5, 115 kDa + smV5, 39 kDa) together with a lower molecular weight band suggestive of a previously characterised Unc-5 cleavage product. **c**, Staged embryos from stages 10–17 showing neural membrane (HRP, magenta) together with endogenous Unc-5 (Unc-5^::smV5^, yellow) localisation during development (Scale bar: 50 µm). **c’**, High magnification of stage 15–16 VNC (Scale bar: 20 µm). **d**, Unc-5^::smV5^ rescues lethality associated with homozygote null mutants (Unc-5^8^) suggesting functionality. **e–i**, Representative traces and violin plots showing Unc-5^::smV5^ does not infer clear neurotransmission defects (Scale Bars: EJCs, 1 nA/30 ms mEJCs, 0.1 nA/500 ms). **j**, Scatter plots showing a modest positive linear correlation between relative Unc-5 intensity and spontaneous *Pr* at individual AZs. **k**, This trend was not observed with brp intensity (*n* = 108 AZs). Violin Plots represent median & IQR. All detailed statistical analysis & genotype details are reported in Supplementary Table 2.

**Figure S4.**
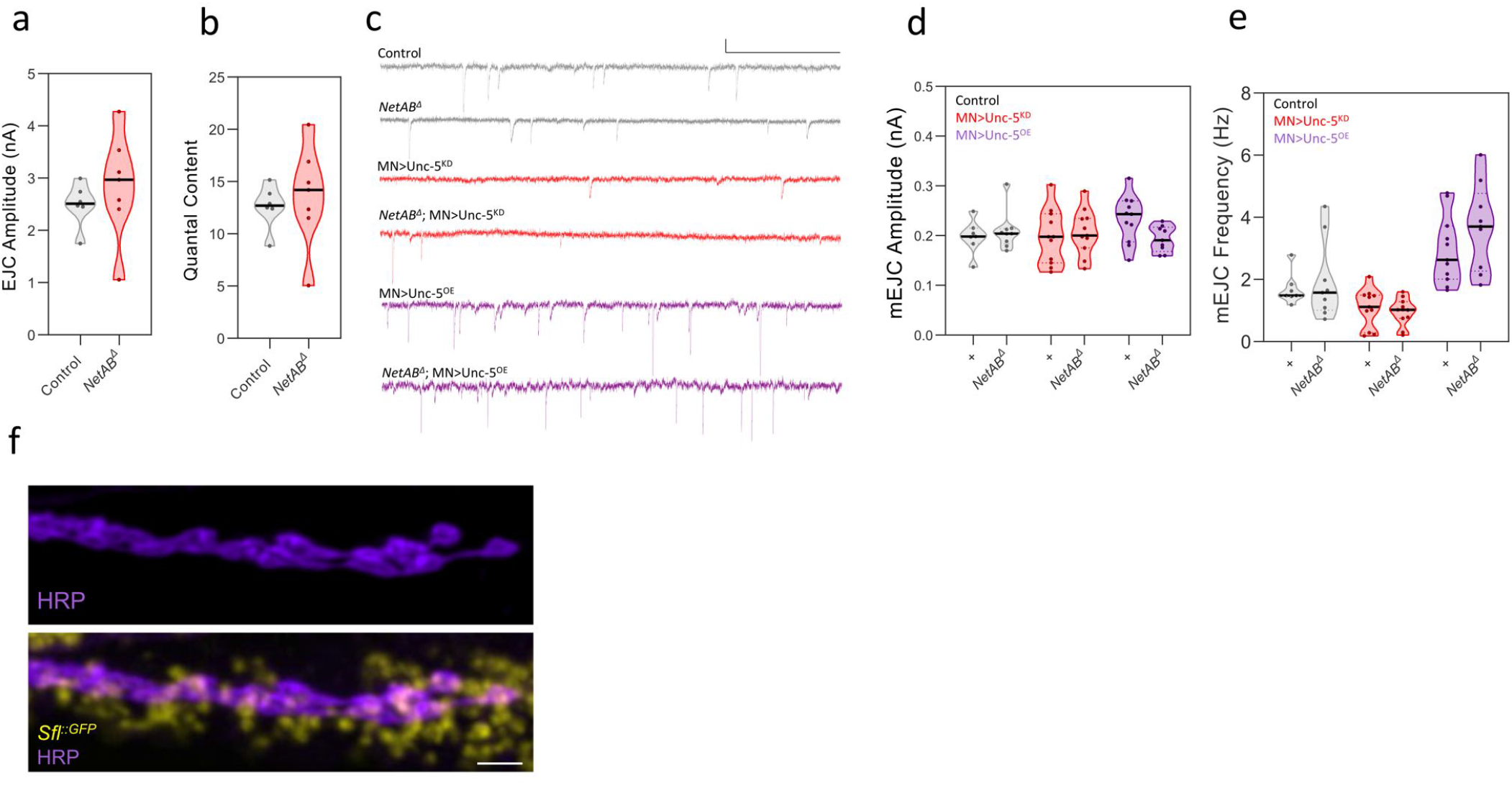
Unc-5 mediated neurotransmission is Netrin independent. **a–b**, No observable change in EJC amplitude or quantal content in animals lacking netrin. **c–e**, No modification to mEJC amplitude or frequency was observed when netrin was deleted in a wildtype background and was not sufficient to modify the penetrance of Unc-5 mediated mEJC frequency modification (Scale bar: 0.1 nA/1 s). **f**, Representative Z-stack image of *sfl*^::GFP^ (yellow) and terminal membrane (HRP, magenta) (Scale bar: 5 µm). Violin Plots represent median & IQR. All detailed statistical analysis & genotype details are reported in Supplementary Table 2.

**Figure S5.**
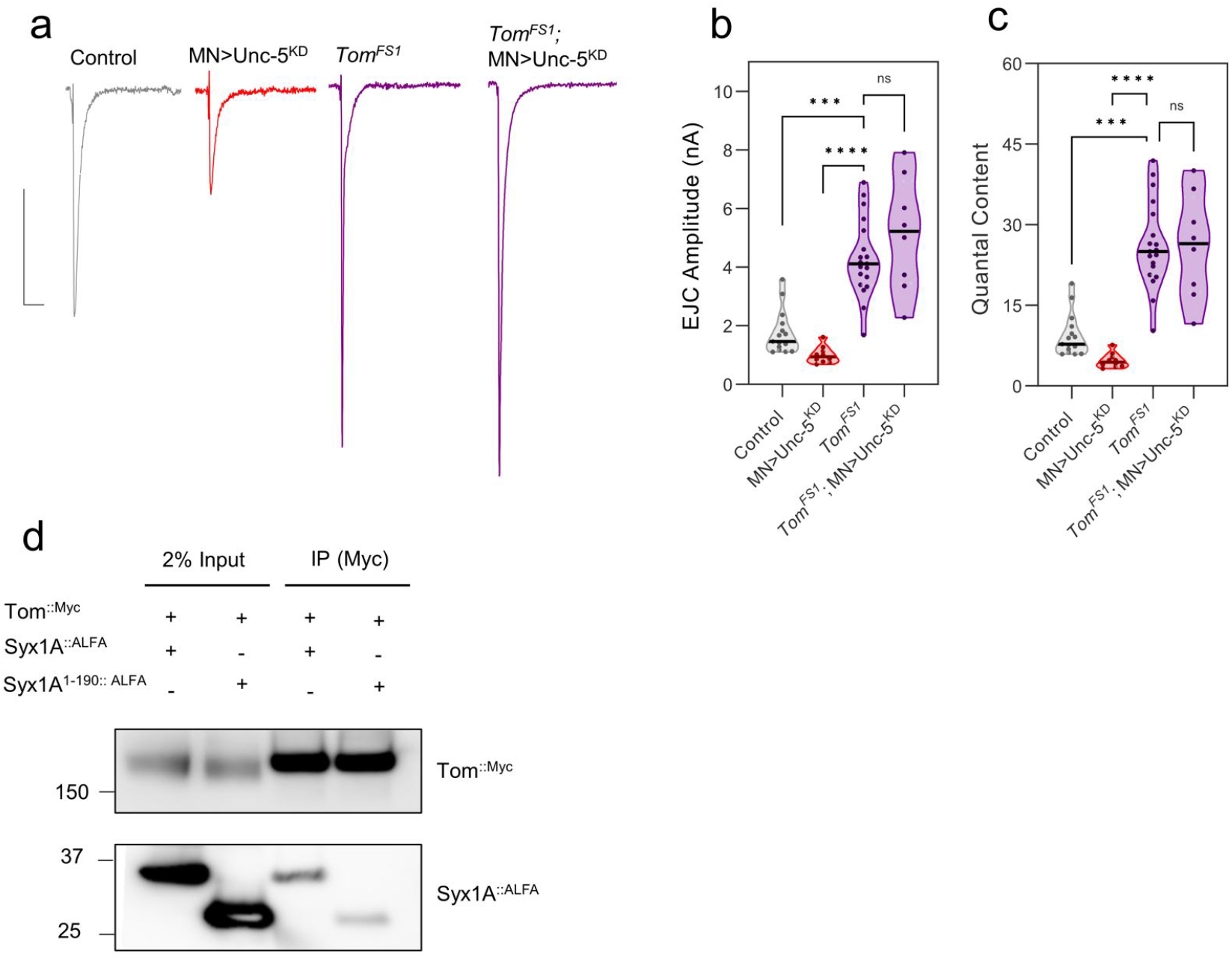
Unc-5 physically interacts with Syx1A and requires the Syx1A C-terminal SNARE domain. **a**, Representative traces displaying EJCs during conditional Unc-5 depletion and Tomosyn deletion (*Tom*^FS1^) (Scale Bars: EJC, 1 nA/30 ms). **b–c**, *Tom*^FS1^ mediated EJC and quantal content potentiation cannot be ameliorated through Unc-5 depletion suggesting, unlike mEJC decline, Unc-5 mediated evoked decline requires Tomosyn. **d**, Tomosyn physically associates with Syx1A and requires the Syx1A C-terminal SNARE domain confirming previously published findings. Violin Plots represent median & IQR. QC was calculated from mEJC amplitudes presented in **Fig. 3e**. All detailed statistical analysis & genotype details are reported in Supplementary Table 2.

**Figure S6.**
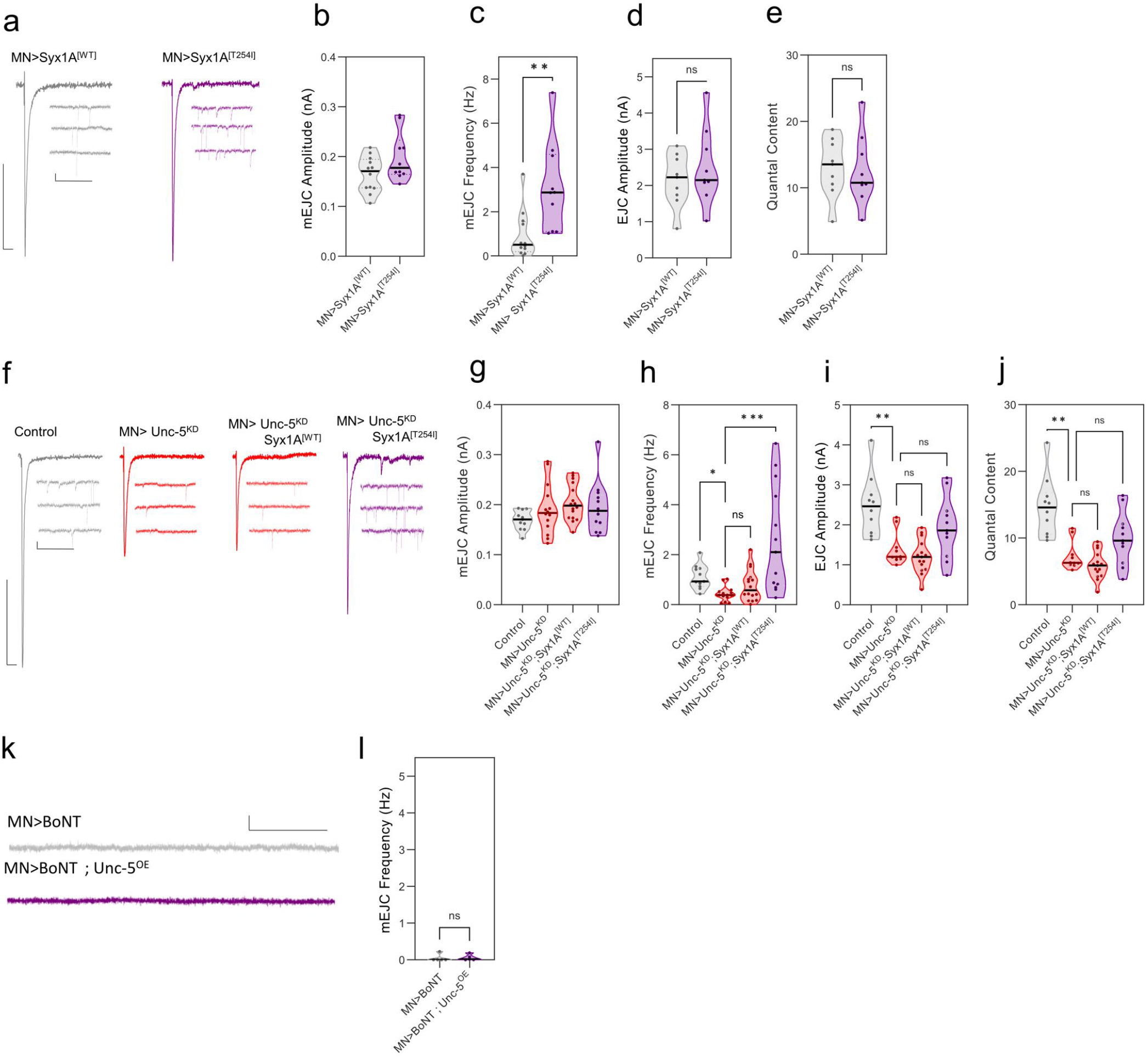
Unc-5 mediated loss of miniature neurotransmission can be rescued by Syx1AT254I. **a**, Representative traces of excitatory glutamatergic currents under conditional motor neuron expression of wildtype Syx1A (Syx1A^WT^) or a hypermorphic mutant known to increase neurotransmitter release (Syx1A^T254I^) (Scale Bars: EJC, 1 nA/30 ms, mEJC, 0.1 nA/500 ms). **b**, Violin plots showing conditional expression of Syx1A^T254I^ did not influence mEJC amplitude. **c**, However, increased mEJC frequency (*n >* 10 recordings per genotype, Student’s *t*-test, *P <* 0.01), **d–e**, and produced no observable effect on EJC amplitude or QC. **f**, Representative traces of excitatory glutamatergic currents under conditional motor neuron depletion of Unc-5 concurrently with transgenic upregulation of Syx1A^WT^ or Syx1A^T254I^ (Scale Bars: EJC, 1 nA/30 ms mEJC, 0.1 nA/500 ms). **g**, Violin plots showing either Syx1A^WT^ or Syx1A^T254I^ did not alter mEJC amplitude during Unc-5 depletion. **h**, However, Syx1A^T254I^ was sufficient to dramatically rescue mEJC frequency (*n >* 11 per genotype, Dunn’s multiple comparisons test, *P <* 0.001) whereas Syx1A^WT^ did not. **ij**, Syx1A^T254I^ produced a partial rescue of EJC amplitude and QC; this however was statistically non-significant when compared to Unc-5 depletion alone. **kl**, Unc-5 unable to increase the frequency of BoNT induced mEJC frequency decline (*P* = 0.85, Scale bar 0.1 nA/500 ms). Violin Plots represent median & IQR. All detailed statistical analysis & genotype details are reported in Supplementary Table 2.

## Supplementary Tables, Main Figures

**Supplementary Table 1.**
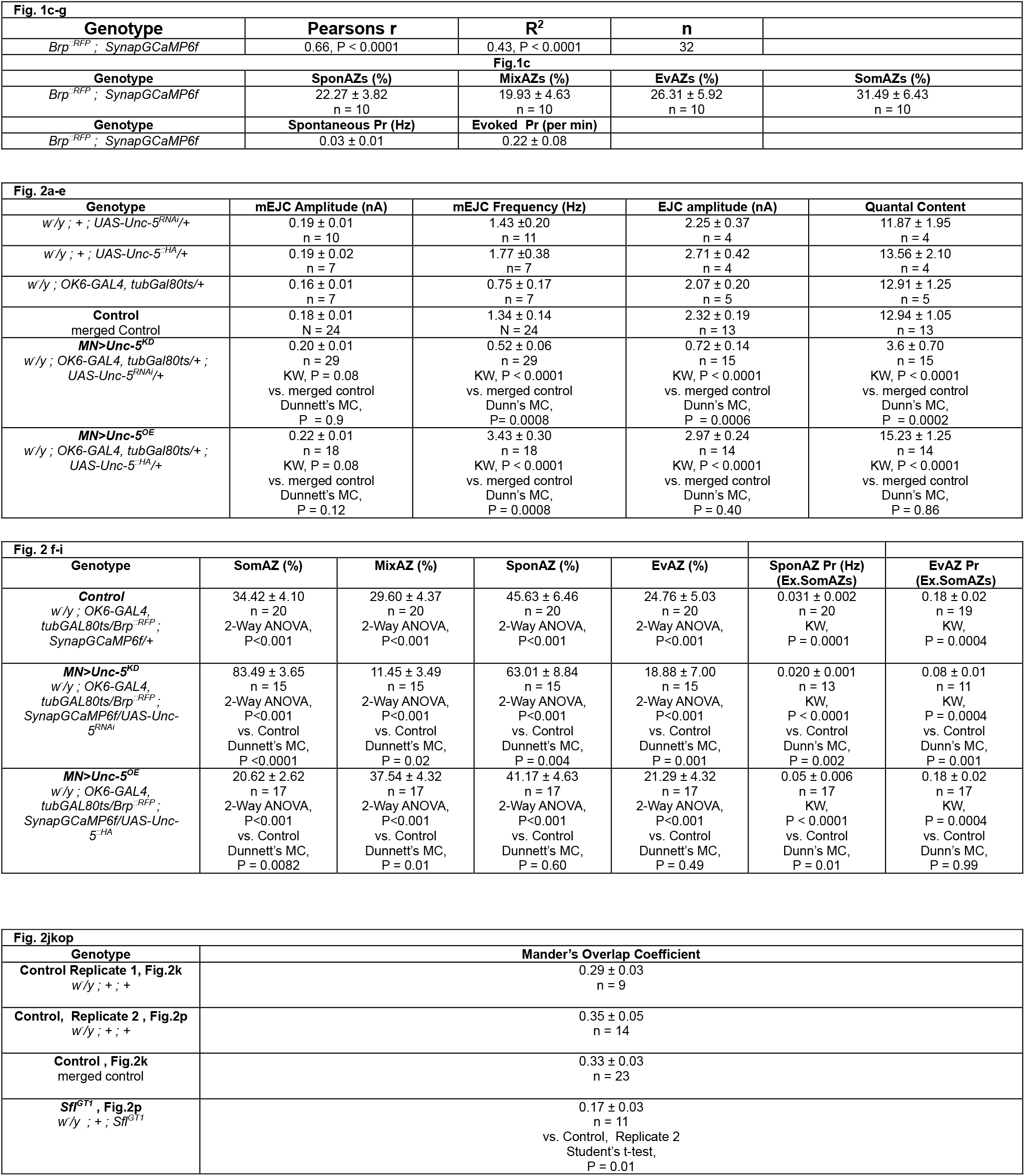
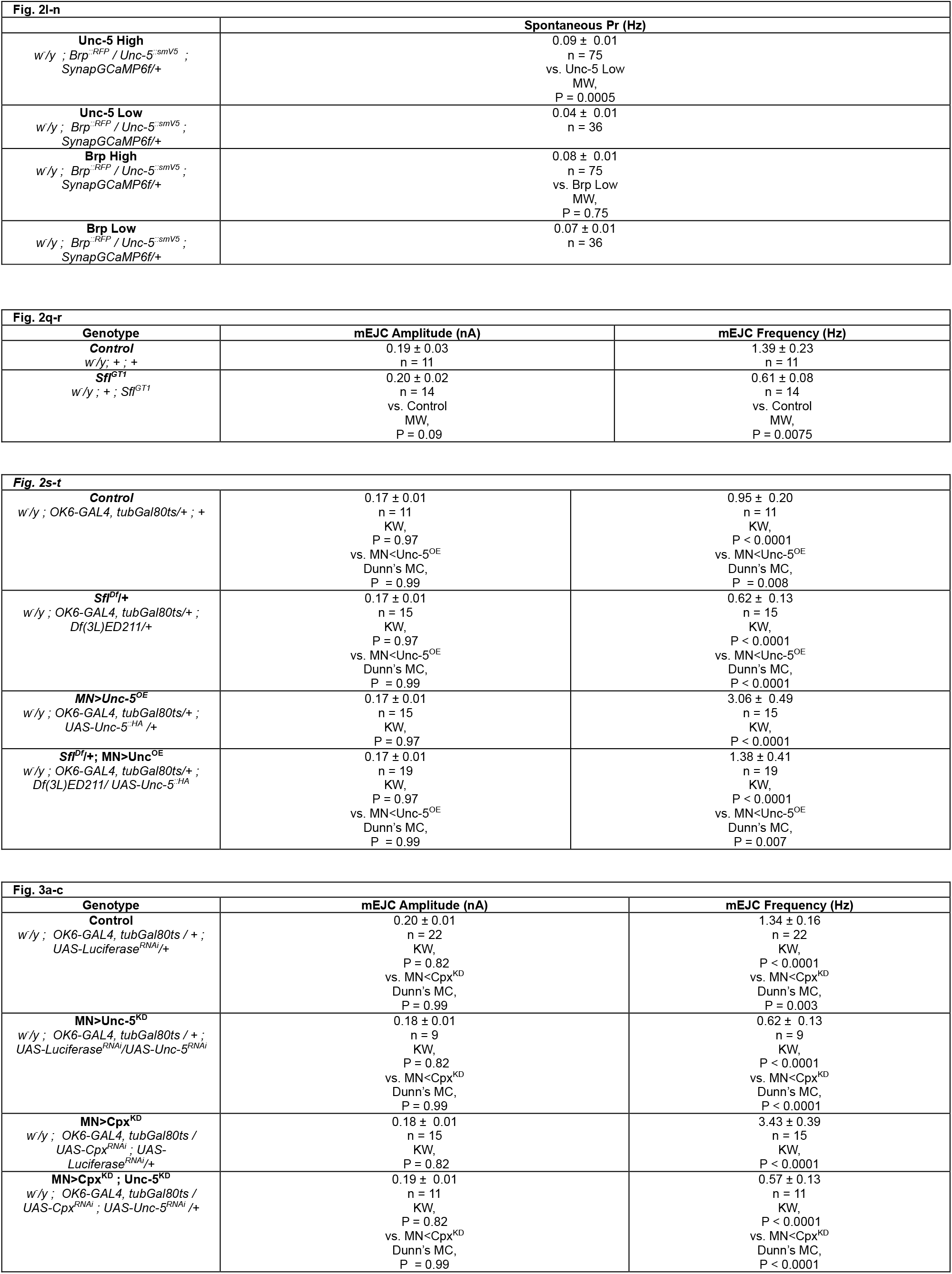
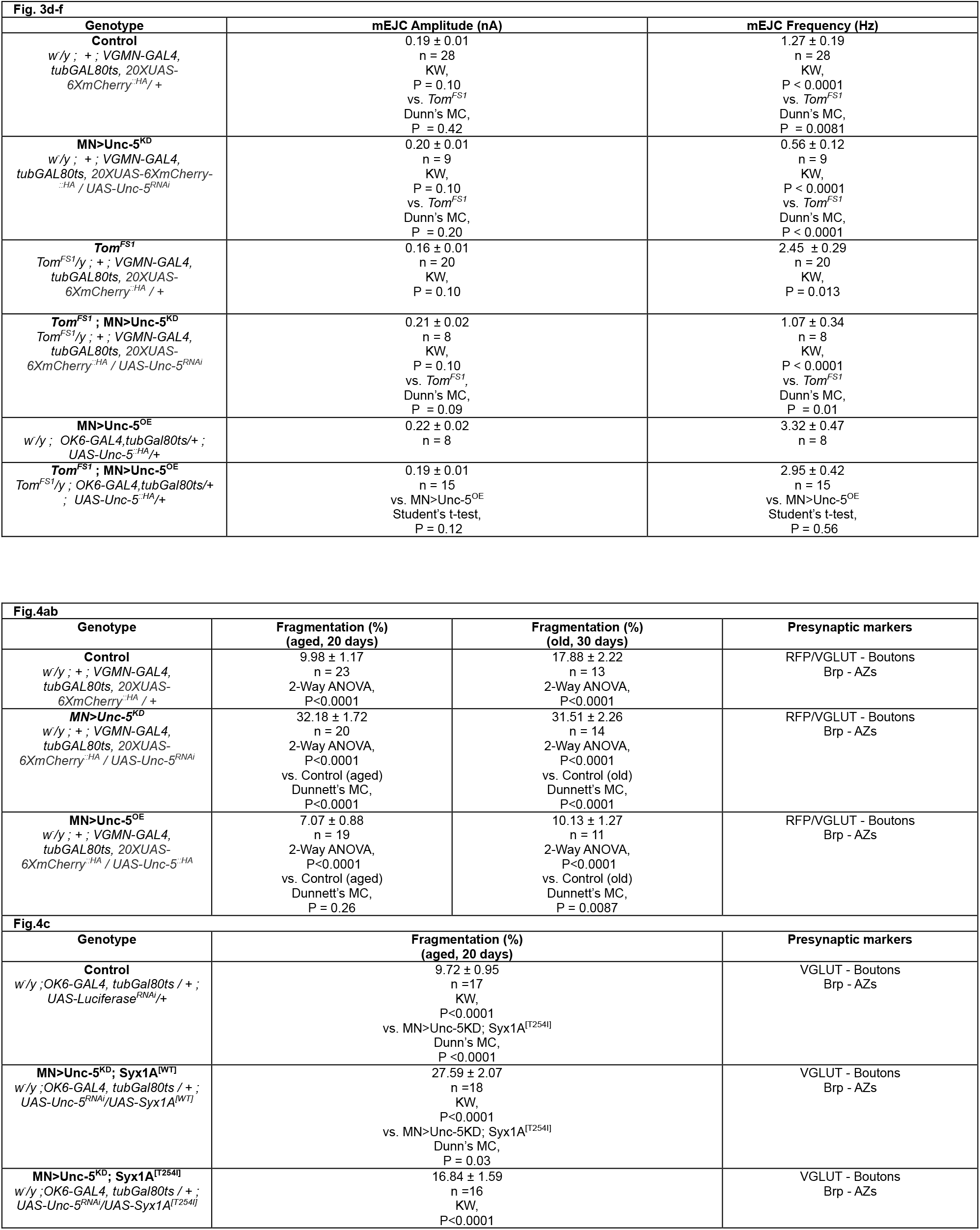
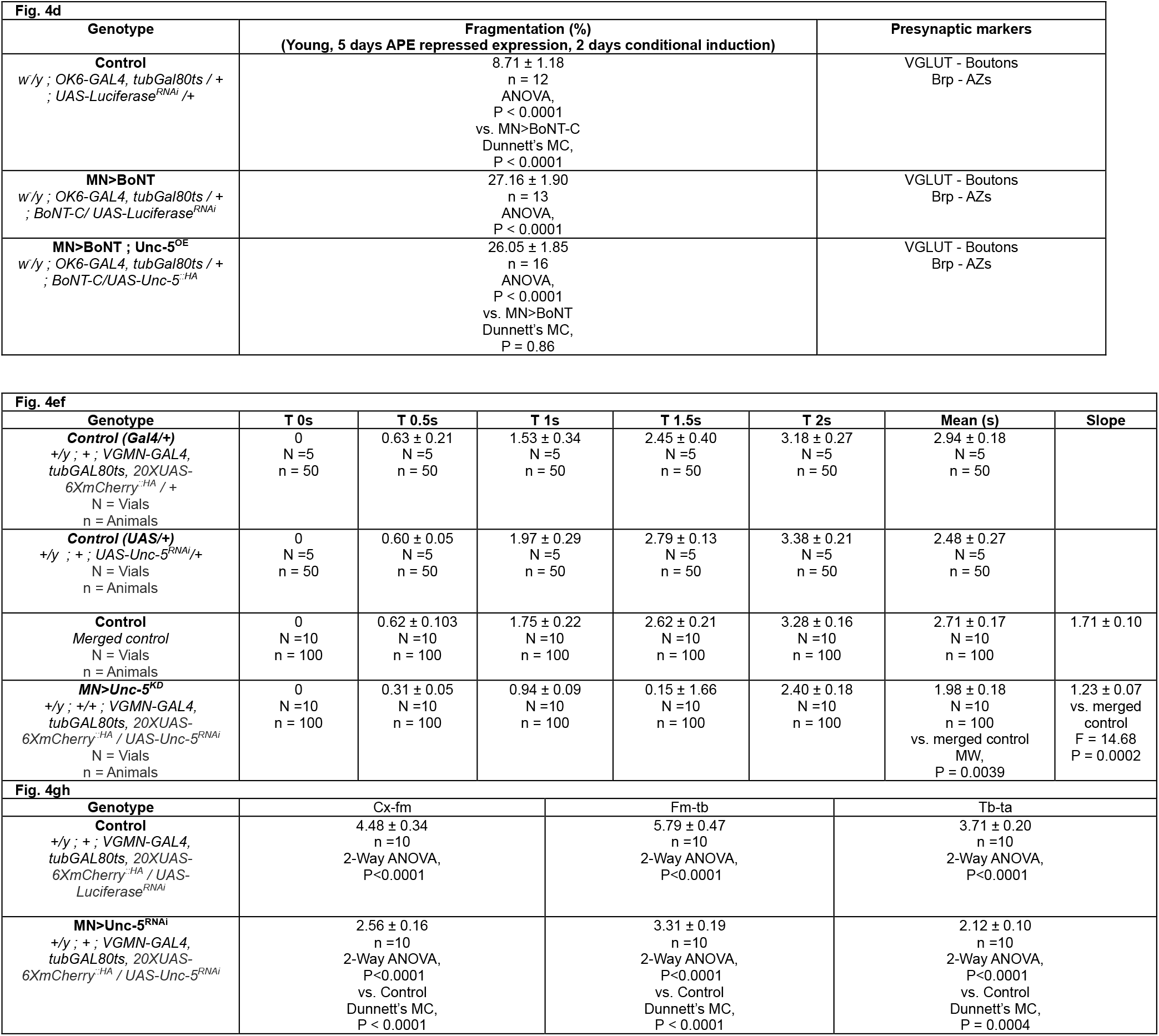

**Supplementary Table 2.**
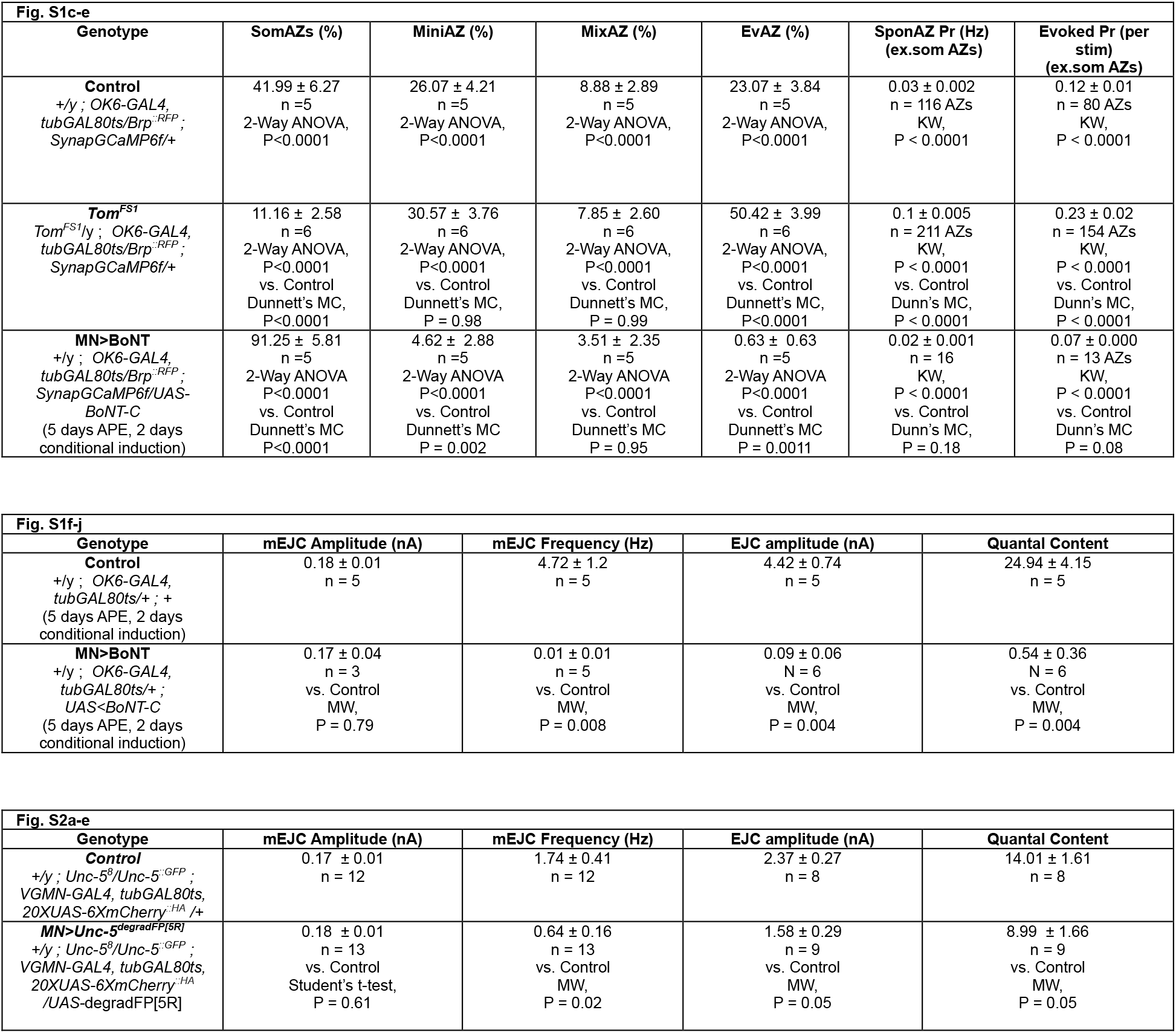
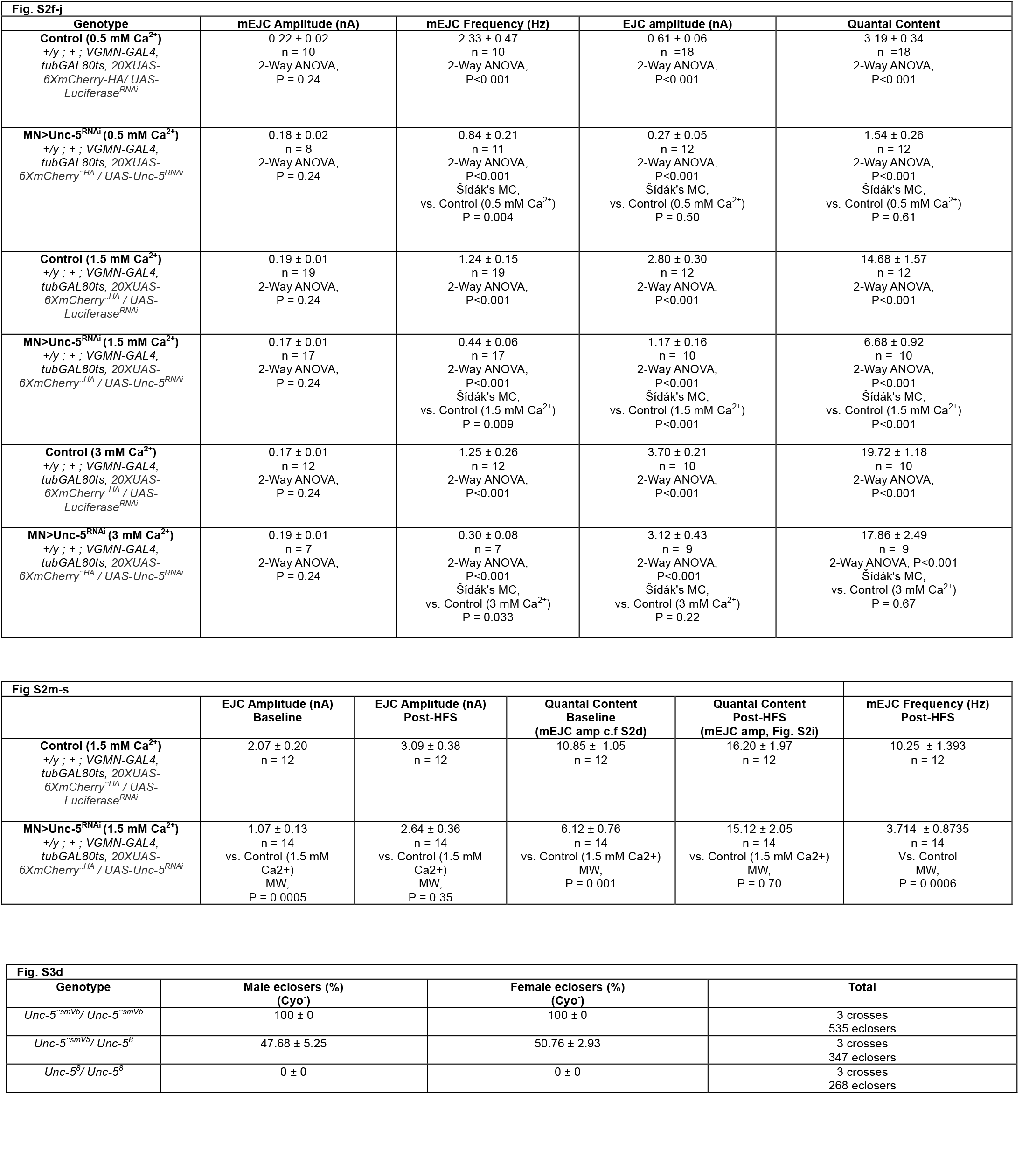
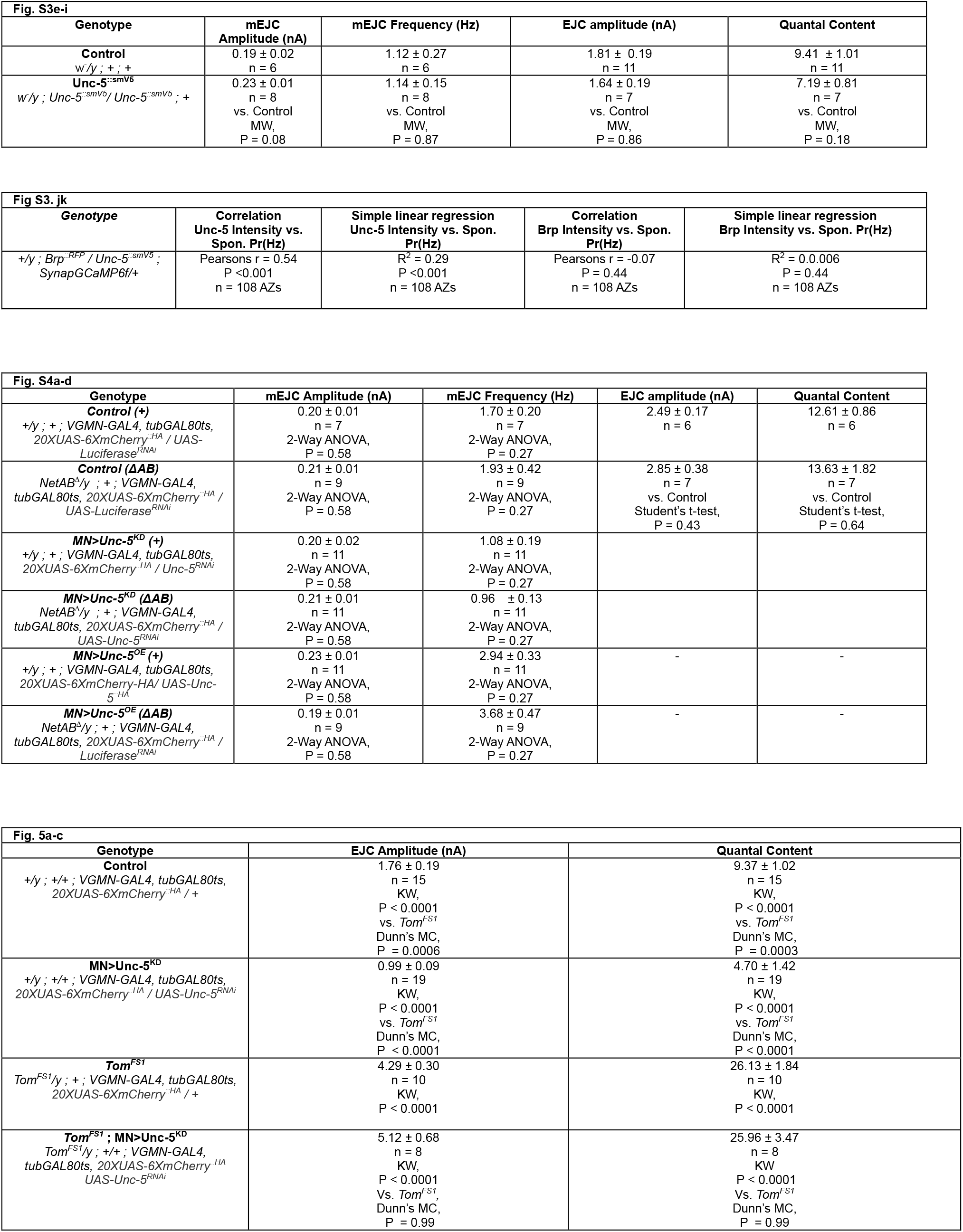
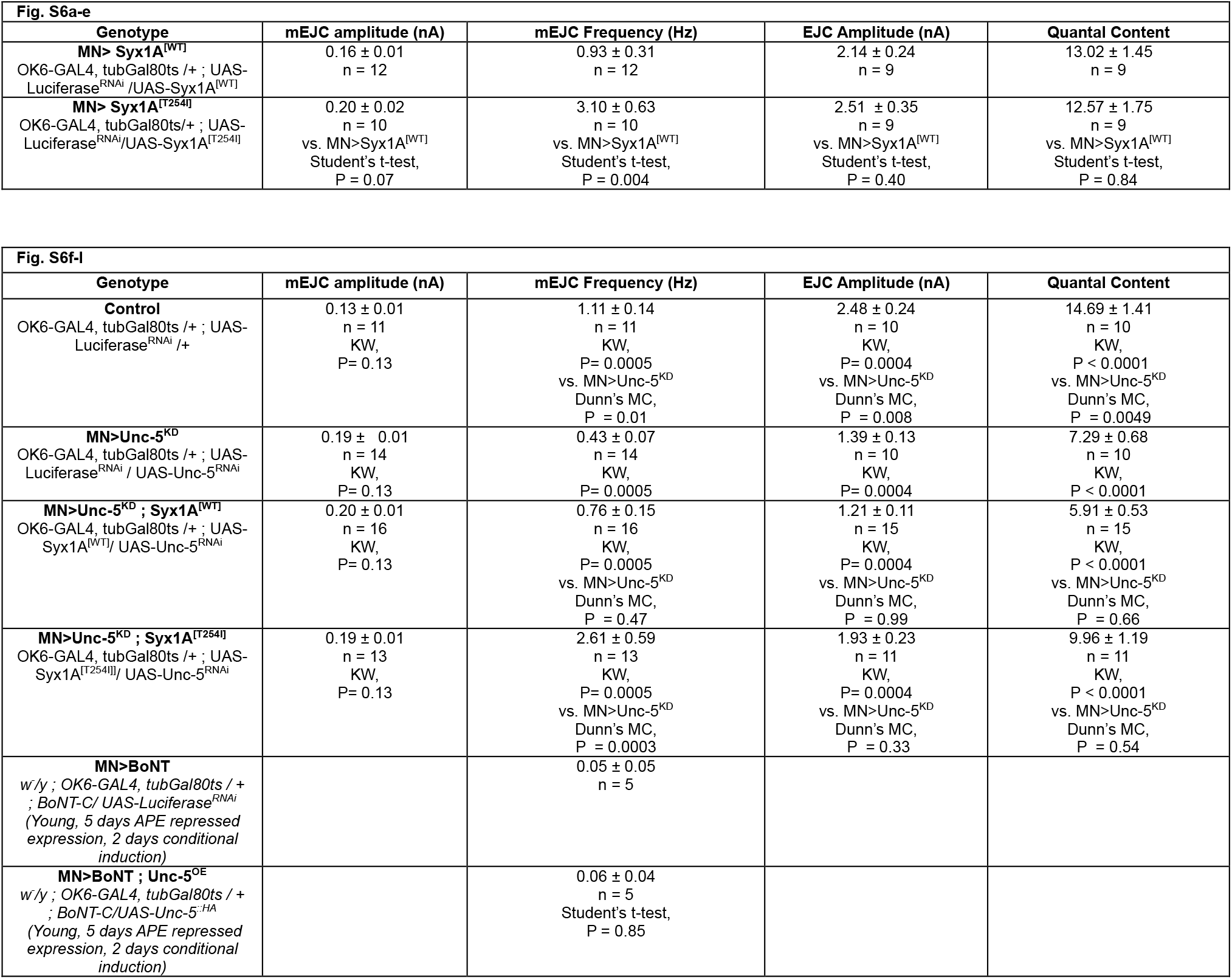
Supplementary Figures.

**Supplementary Table 3.**
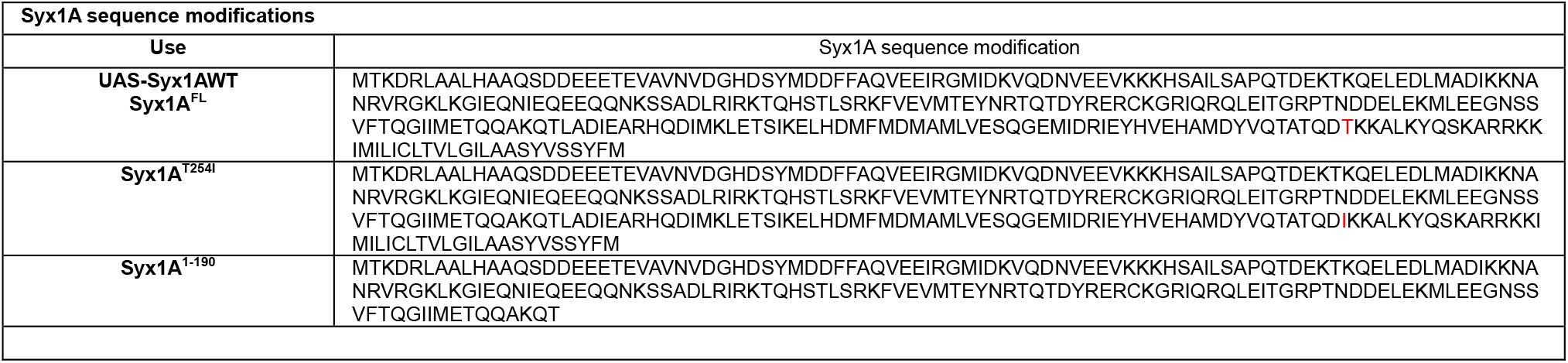

## Bibliography

1. Fatt, P. & Katz, B. Some observations on biological noise. Nature 166, 597–598 (1950).

2. Fatt, P. & Katz, B. Spontaneous subthreshold activity at motor nerve endings. J. Physiol. 117, 109 (1952).

3. Katz, B. & Miledi, R. The measurement of synaptic delay, and the time course of acetylcholine release at the neuromuscular junction. Proc. R. Soc. Lond. B 161, 483–495 (1965).

4. Heuser, J. E. & Reese, T. Evidence for recycling of synaptic vesicle membrane during transmitter release at the frog neuromuscular junction. J. Cell Biol. 57, 315–344 (1973).

5. Ceccarelli, B. & Hurlbut, W. P. Vesicle hypothesis of the release of quanta of acetylcholine. Physiol. Rev. 60, 396–441 (1980).

6. Südhof, T. C. The synaptic vesicle cycle. Annu. Rev. Neurosci. 27, 509–547 (2004).

7. Kavalali, E. T. The mechanisms and functions of spontaneous neurotransmitter release. Nat. Rev. Neurosci. 16, 5–16 (2015).

8. Del Castillo, J. & Katz, B. Quantal components of the end-plate potential. J. Physiol. 124, 560 (1954).

9. Martin, A. R. Quantal nature of synaptic transmission. Physiol. Rev. (1966).

10. Redman, S. Quantal analysis of synaptic potentials in neurons of the central nervous system. Physiol. Rev. 70, 165–198 (1990).

11. Faisal, A. A., Selen, L. P. & Wolpert, D. M. Noise in the nervous system. Nat. Rev. Neurosci. 9, 292–303 (2008).

12. Tyler, W. J. & Pozzo-Miller, L. Miniature synaptic transmission and BDNF modulate dendritic spine growth and form in rat CA1 neurones. J. Physiol. 553, 497–509 (2003).

13. Murase, S., Mosser, E. & Schuman, E. M. Depolarization drives *β*-catenin into neuronal spines promoting changes in synaptic structure and function. Neuron 35, 91–105 (2002).

14. Chao, H. T., Zoghbi, H. Y. & Rosenmund, C. MeCP2 controls excitatory synaptic strength by regulating glutamatergic synapse number. Neuron 56, 58–65 (2007).

15. Tran, T. S. et al. Secreted semaphorins control spine distribution and morphogenesis in the postnatal CNS. Nature 462, 1065–1069 (2009).

16. Shao, L. X. et al. Psilocybin induces rapid and persistent growth of dendritic spines in frontal cortex in vivo. Neuron 109, 2535–2544 (2021).

17. El-Husseini, A. E. D. et al. PSD-95 involvement in maturation of excitatory synapses. Science 290, 1364–1368 (2000).

18. Mackenzie, P. J. et al. Vesicle number does not predict postsynaptic measures of miniature synaptic activity frequency in cultured cortical neurons. Neuroscience 98, 1–7 (2000).

19. Van Roessel, P. et al. Independent regulation of synaptic size and activity by the anaphase-promoting complex. Cell 119, 707–718 (2004).

20. Collins, C. A. et al. Highwire restrains synaptic growth by attenuating a MAP kinase signal. Neuron 51, 57–69 (2006).

21. McCabe, B. D. et al. Highwire regulates presynaptic BMP signaling essential for synaptic growth. Neuron 41, 891–905 (2004).

22. Ramirez, D. M. et al. Vti1a identifies a vesicle pool that preferentially recycles at rest and maintains spontaneous neurotransmission. Neuron 73, 121–134 (2012).

23. Sara, Y. et al. An isolated pool of vesicles recycles at rest and drives spontaneous neurotransmission. Neuron 45, 563–573 (2005).

24. Chung, C. et al. Acute dynamin inhibition dissects synaptic vesicle recycling pathways that drive spontaneous and evoked neurotransmission. J. Neurosci. 30, 1363–1376 (2010).

25. Peled, E. S. & Isacoff, E. Y. Optical quantal analysis of synaptic transmission in wild-type and rab3-mutant *Drosophila* motor axons. Nat. Neurosci. 14, 519–526 (2011).

26. Melom, J. E., Akbergenova, Y., Gavornik, J. P. & Littleton, J. T. Spontaneous and evoked release are independently regulated at individual active zones. J. Neurosci. 33, 17253–17263 (2013).

27. Newman, Z. L. et al. Input-specific plasticity and homeostasis at the *Drosophila* larval neuromuscular junction. Neuron 93, 1388–1404 (2017).

28. Akbergenova, Y. et al. Characterization of developmental and molecular factors underlying release heterogeneity at *Drosophila* synapses. eLife 7, e38268 (2018).

29. Newman, Z. L. et al. Determinants of synapse diversity revealed by super-resolution quantal transmission and active zone imaging. Nat. Commun. 13, 229 (2022).

30. Grasskamp, A. T. et al. Spontaneous neurotransmission at evocable synapses predicts their responsiveness to action potentials. Front. Cell. Neurosci. 17, 1129417 (2023).

31. Wang, C. S., Monteggia, L. M. & Kavalali, E. T. Spatially non-overlapping Ca2+ signals drive distinct forms of neurotransmission. Cell Rep. 42 (2023).

32. Horvath, P. M. et al. Spontaneous and evoked neurotransmission are partially segregated at inhibitory synapses. eLife 9, e52852 (2020).

33. Bal, M. et al. Reelin mobilizes a VAMP7-dependent synaptic vesicle pool and selectively augments spontaneous neurotransmission. Neuron 80, 934–946 (2013).

34. Saitoe, M. et al. Absence of junctional glutamate receptor clusters in *Drosophila* mutants lacking spontaneous transmitter release. Science 293, 514–517 (2001).

35. Choi, B. J. et al. Miniature neurotransmission regulates *Drosophila* synaptic structural maturation. Neuron 82, 618–634 (2014).

36. Andreae, L. C. & Burrone, J. Spontaneous neurotransmitter release shapes dendritic arbors via long-range activation of NMDA receptors. Cell Rep. 10, 873–882 (2015).

37. Sutton, M. A. et al. Miniature neurotransmission stabilizes synaptic function via tonic suppression of local dendritic protein synthesis. Cell 125, 785–799 (2006).

38. Banerjee, S. et al. Miniature neurotransmission is required to maintain *Drosophila* synaptic structures during ageing. Nat. Commun. 12, 4399 (2021).

39. Banerjee, S. et al. Trio preserves motor synapses and prolongs motor ability during aging. Cell Rep. 43 (2024).

40. Südhof, T. C. The presynaptic active zone. Neuron 75, 11–25 (2012).

41. Bindels, D. S. et al. mScarlet: a bright monomeric red fluorescent protein for cellular imaging. Nat. Methods 14, 53–56 (2017).

42. Wagh, D. A. et al. Bruchpilot, a protein with homology to ELKS/CAST, is required for structural integrity and function of synaptic active zones in *Drosophila*. Neuron 49, 833–844 (2006).

43. Han, Y. et al. Botulinum neurotoxin accurately separates tonic vs. phasic transmission and reveals heterosynaptic plasticity rules in *Drosophila*. eLife 11, e77924 (2022).

44. Sauvola, C. W. et al. The decoy SNARE Tomosyn sets tonic versus phasic release properties and is required for homeostatic synaptic plasticity. eLife 10, e72841 (2021).

45. Boyer, N. P. & Gupton, S. L. Revisiting netrin-1: one who guides (axons). Front. Cell. Neurosci. 12, 221 (2018).

46. Li, Q. et al. The role of UNC5C in Alzheimer’s disease. Ann. Transl. Med. 6, 178 (2018).

47. Zhou, Z. et al. A systematic analysis of the role of UNC-5 netrin receptor a (UNC5A) in human cancers. Biomolecules 12, 1826 (2022).

48. Caussinus, E. & Affolter, M. deGradFP: a system to knockdown GFP-tagged proteins. Drosophila: Methods and Protocols 177–187 (2016).

49. Källstig, E. et al. Highly frequent undesired insertional mutagenesis during *Drosophila* genome editing. bioRxiv (2025).

50. Viswanathan, S. et al. High-performance probes for light and electron microscopy. Nat. Methods 12, 568–576 (2015).

51. Keleman, K. & Dickson, B. J. Short- and long-range repulsion by the *Drosophila* Unc-5 netrin receptor. Neuron 32, 605–617 (2001).

52. Leonardo, E. D. et al. Vertebrate homologues of *C. elegans* UNC-5 are candidate netrin receptors. Nature 386, 833–838 (1997).

53. Grandin, M. et al. Structural decoding of the Netrin-1/UNC5 interaction and its therapeutical implications in cancers. Cancer Cell 29, 173–185 (2016).

54. Newquist, G. et al. Control of male and female fertility by the netrin axon guidance genes. PLoS One 8, e72524 (2013).

55. Akkermans, O. et al. GPC3-Unc5 receptor complex structure and role in cell migration. Cell 185, 3931–3949 (2022).

56. Priest, J. M. et al. Structural insights into the formation of repulsive netrin guidance complexes. Sci. Adv. 10, eadj8083 (2024).

57. Habuchi, O. Diversity and functions of glycosaminoglycan sulfotransferases. Biochim. Biophys. Acta 1474, 115–127 (2000).

58. Lukacsovich, T. et al. Dual-tagging gene trap of novel genes in *Drosophila melanogaster*. Genetics 157, 727–742 (2001).

59. Bellen, H. J. et al. The BDGP gene disruption project: single transposon insertions associated with 40% of *Drosophila* genes. Genetics 167, 761–781 (2004).

60. Rizo, J. & Xu, J. The synaptic vesicle release machinery. Annu. Rev. Biophys. 44, 339–367 (2015).

61. Huntwork, S. & Littleton, J. T. A complexin fusion clamp regulates spontaneous neurotransmitter release and synaptic growth. Nat. Neurosci. 10, 1235–1237 (2007).

62. Martínez-Mármol, R. et al. Syntaxin-1 is necessary for UNC5A-C/Netrin-1-dependent macropinocytosis and chemorepulsion. Front. Mol. Neurosci. 16, 1253954 (2023).

63. Mahadik, S. S. & Lundquist, E. A. TOM-1/tomosyn acts with the UNC-6/netrin receptor UNC-5 to inhibit growth cone protrusion in *Caenorhabditis elegans*. Development 150, dev201031 (2023).

64. Bennett, M. K., Calakos, N. & Scheller, R. H. Syntaxin: a synaptic protein implicated in docking of synaptic vesicles at presynaptic active zones. Science 257, 255–259 (1992).

65. Fujita, Y. et al. Tomosyn: a syntaxin-1-binding protein that forms a novel complex in the neurotransmitter release process. Neuron 20, 905–915 (1998).

66. Lagow, R. D. et al. Modification of a hydrophobic layer by a point mutation in syntaxin 1A regulates the rate of synaptic vesicle fusion. PLoS Biol. 5, e72 (2007).

67. Vernon, S. W., Goodchild, J. & Baines, R. A. The VAChTY 49N mutation provides insecticide-resistance but perturbs evoked cholinergic neurotransmission in *Drosophila*. PLoS One 13, e0203852 (2018).

68. Mathis, A. et al. DeepLabCut: markerless pose estimation of user-defined body parts with deep learning. Nat. Neurosci. 21, 1281–1289 (2018).

69. Lobato-Rios, V. et al. NeuroMechFly, a neuromechanical model of adult *Drosophila melanogaster*. Nat. Methods 19, 620–627 (2022).

70. Aberle, H. et al. wishful thinking encodes a BMP type II receptor that regulates synaptic growth in *Drosophila*. Neuron 33, 545–558 (2002).

71. Ruchti, E. & McCabe, B. D. Tools for simultaneous synchronous temporal control of *Drosophila* gene expression in discrete cells. bioRxiv (2026).

72. Shearin, H. K. et al. Hexameric GFP and mCherry reporters for the *Drosophila* GAL4, Q, and LexA transcription systems. Genetics 196, 951–960 (2014).

73. McGuire, S. E. et al. Spatiotemporal rescue of memory dysfunction in *Drosophila*. Science 302, 1765–1768 (2003).

74. Labrador, J. P. et al. The homeobox transcription factor even-skipped regulates netrin-receptor expression to control dorsal motor-axon projections in *Drosophila*. Curr. Biol. 15, 1413–1419 (2005).

75. Nagarkar-Jaiswal, S. et al. A library of MiMICs allows tagging of genes and reversible, spatial and temporal knockdown of proteins in *Drosophila*. eLife 4, e05338 (2015).

76. Nern, A. et al. Multiple new site-specific recombinases for use in manipulating animal genomes. Proc. Natl. Acad. Sci. USA 108, 14198–14203 (2011).

77. Daniel, K. et al. Conditional control of fluorescent protein degradation by an auxin-dependent nanobody. Nat. Commun. 9, 3297 (2018).

78. Stapleton, M. et al. A *Drosophila* full-length cDNA resource. Genome Biol. 3, research0080–1 (2002).

79. Coker, J. A. et al. FAS2FURIOUS: moderate-throughput secreted expression of difficult recombinant proteins in *Drosophila* S2 cells. Front. Bioeng. Biotechnol. 10, 871933 (2022).

80. Bashaw, G. J. Visualizing axons in the *Drosophila* central nervous system using immunohistochemistry and immunofluorescence. Cold Spring Harb. Protoc. 2010 (2010).

81. Dervinis, M. & Major, G. A novel method for reliably measuring miniature and spontaneous postsynaptic events in whole-cell patch clamp recordings. Front. Cell. Neurosci. 19, 1598016 (2025).

82. Spierer, A. N. et al. FreeClimber: automated quantification of climbing performance in *Drosophila*. J. Exp. Biol. 224, jeb229377 (2021).

